# ALPARC: Artificial Languages with Phonological and Acoustic Rhythmicity Controls

**DOI:** 10.1101/2024.05.24.595268

**Authors:** Lorenzo Titone, Nikola Milosevic, Lars Meyer

## Abstract

Infants and adults show the remarkable ability to learn from statistical regularities in the environment. Seminal studies on statistical learning in language acquisition suggested that transitional probabilities between syllables are decisive for word learning. Yet, recent work cautioned that acoustic and phonological regularities confound transitional probabilities, compromising interpretability. Furthermore, prior linguistic background can impact the learning of a new (artificial) language. To control for such confounds, we developed an open-source Python toolbox that generates Artificial Languages with Phonological and Acoustic Rhythmicity Controls (ALPARC). First, we explain all functionalities of ALPARC and provide a step-by-step guide. Then, we demonstrate how ALPARC generates syllable streams encompassing pseudowords that are tailored to critical statistics of real languages. Our results show that ALPARC streams attain stationary transitional probability distributions and reduce acoustic and phonological confounds relative to stimuli used in prior studies. We conclude that ALPARC is a useful tool to overcome current uncertainties in future SL studies.

## 1. Introduction

Statistical learning (SL) is the ability to extract regularities from the environment. Among its role for several cognitive functions, SL is pivotal for speech segmentation and language acquisition (Bogaerts et al., 2020; Frost et al., 2019). For instance, statistical regularities can be used to infer lexical boundaries that are rarely marked acoustically, since transitional probabilities (TPs) between syllables are typically higher within than across words. Seminal research on SL in language acquisition used artificial language learning paradigms to show that infants and adults can acquire a lexicon of nonsense “pseudowords” marked by low-TP boundaries (e.g., Saffran et al., 1996). After exposure to syllable streams with statistical regularities, SL can be assessed by reflection-based measures, such as two-alternative forced-choice tasks, where pseudowords are endorsed as more familiar than chunks that violate the TPs (see Batterink et al., 2015; Christiansen, 2019, for discussion).

Several electrophysiological studies used frequency-tagging paradigms to investigate neural tracking of periodic TP patterns as an index of linguistic SL (Assaneo et al., 2019; Batterink & Paller, 2017; Buiatti et al., 2009; Cunillera et al., 2006, 2009; Getz et al., 2018; Henin et al., 2021; McNealy et al., 2006; Ordin et al., 2020). In this paradigm, syllables are presented at a fixed rate (e.g., 3.3 Hz), while pseudowords emerge at a different rate (e.g., 1.1 Hz) in TP-structured streams, but not in shuffled streams. Most of these studies found decreased neural tracking at the syllable rate (physically present in the stimulus) and increased tracking at the pseudoword rate (putatively absent from the stimulus) in TP-structured relative to TP-uniform streams (see also Sjuls et al., 2024 for a review). While these findings suggest sensitivity to TPs, or perceptual chunking of pseudowords (see Batterink & Paller, 2017; Perruchet & Pacton, 2006; Saffran & Kirkham, 2018; Siegelman et al., 2019; Thiessen, 2017 for discussion), recent work raised concerns about the interpretability of this work since acoustic regularities and phonotactics confound the TP rate (Pinto et al., 2022; van der Wulp, 2021).

Pinto et al. (2022) pointed out that syllables occurring at fixed positions in TP-structured streams impose an acoustic peak at the pseudoword rate in the envelope spectra of TP-structured streams, which is absent in TP-uniform streams with randomly shuffled syllables (Luo & Ding, 2020; Pinto et al., 2022). In order to match this acoustic peak at baseline, Pinto et al. (2022) designed a TP-uniform position-controlled stream, forcing syllables to occupy the same ordinal position. Moreover, they found stronger tracking at the pseudoword rate in the TP-structured condition than in the baseline. While this acoustic control strengthens neural inference, several other factors may still play a role in pseudoword learning, beyond TPs. In fact, one caveat of linguistic SL is that learners may infer lexical boundaries also based on acoustic, phonological, and phonotactic cues (Luo & Ding, 2020; Onnis et al., 2005; Pinto et al., 2022; Seidenberg et al., 2002; van der Wulp, 2021; Varela et al., 2024). Indeed, neurons in the human superior temporal gyrus display selective tuning to phonemes and phonological features (Mesgarani et al., 2014). Accordingly, studies have suggested that cortical tracking of speech might reflect phoneme-level processing in addition to statistical regularities or acoustic cues (Chen et al., 2020; Daube et al., 2019; Di Liberto et al., 2015).

Phonotactic rules in a given language can bias word segmentation. For instance, neighboring syllables with the same place of articulation are more likely to occur across word boundaries than within words. The adherence to this phonotactic rule—also known as the obligatory contour principle (OCP)—was shown to influence word segmentation (Boll-Avetisyan & Kager, 2014) and to partially account for neural tracking at the pseudoword rate in a recent reanalysis of an artificial language learning study (Batterink & Paller, 2017; van der Wulp, 2021). Moreover, previous studies using coarticulation or stress in combination with statistical linguistic information showed that infants rely on prosodic and phonotactic cues (Johnson & Jusczyk, 2001; Jusczyk et al., 1999; Thiessen & Saffran, 2003). Building on this work, studies have shown the role of familiar prosodic cues for language learning (Kuuluvainen et al., 2025; Shukla et al., 2007), as well as evidence for neural tracking of both prosody and TPs with frequency-tagging (Cunillera et al., 2006; Elmer et al., 2021). In addition to acoustic, phonological, prosodic, and statistical regularities, linguistic entrenchment with a known language biases the learning of a new (artificial) language (Elazar et al., 2022; Siegelman et al., 2018; Varela et al., 2024). To address this issue, prior work relied on experts’ knowledge and linguistic corpora to control the similarity between artificial and real languages by excluding syllables with high frequency of use (Kiai & Melloni, 2021) or eluding high positional similarity to real words (Fernandes et al., 2009; Varela et al., 2024).

We introduce here an open-source tool to improve stimulus control in (linguistic) SL studies. The tool generates *Artificial Languages with Phonological and Acoustic Rhythmicity Controls* (ALPARC). ALPARC is optimized to make frequency-tagged syllable streams with regular TP distributions. In ALPARC, artificial languages are defined as sets of pseudowords. ALPARC builds these sets strategically to prevent phonological and linguistic cues from biasing the TPs between pseudowords. These sets are then rendered in TP-structured and TP-uniform streams, which can be (externally) synthesized or recorded, and further edited using ad-hoc duration and pitch manipulations for frequency-tagging paradigms. In addition, ALPARC introduces a new Phonological Rhythmicity Index (PRI) diagnosing the presence of phonological regularities at the TP rate in such TP-uniform streams of syllables.

We first highlight the current scope of ALPARC with three independent and interdependent modules. ALPARC—1 builds artificial languages (of arbitrary size) from a list of pseudoword candidates (with a fixed syllable structure). ALPARC—2 renders these artificial languages as TP-structured and TP-uniform streams, contrasting surprisal at word boundaries. ALPARC—3 diagnoses a PRI that reflects phonological rhythmicity confounds at the pseudoword rate in frequency-tagged speech streams. Afterwards, we provide a step-by-step guide to simulate artificial lexicons of variable size, which are rendered in TP-uniform syllable streams of arbitrary length with calibrated statistical, phonological, and acoustic rhythmicity at a fixed rate. We evaluate ALPARC showing that it reduces acoustic and phonological rhythmicity confounds at the rate of interest and enhances TP stationarity relative to benchmarking artificial languages and naïve randomness methods. Finally, we discuss the current limitations and potential future directions of every ALPARC module.

## 2. Materials and Equipment

ALPARC is an open-source tool with a modular architecture. It can be used as a dataset to download our pre-computed artificial languages, or as a library to (1) generate new artificial languages, (2) render them as TP-structured and TP-uniform streams, and (3) evaluate phonological rhythmicity confounds therein. ALPARC achieves these goals in five steps, optimized for linguistic SL studies with frequency-tagging. In parallel to assuming a linear workflow, the architecture of ALPARC can be divided into three modules, according to their functional domains (Figure 1). Within this scope, ALPARC has a wide array of parameter settings (Table 1). Beyond the present scope, the three modules of ALPARC might serve as independent base tools for future SL research across domains. All materials, data, and code are available at: https://github.com/milosen/alparc.

**Figure 1.**
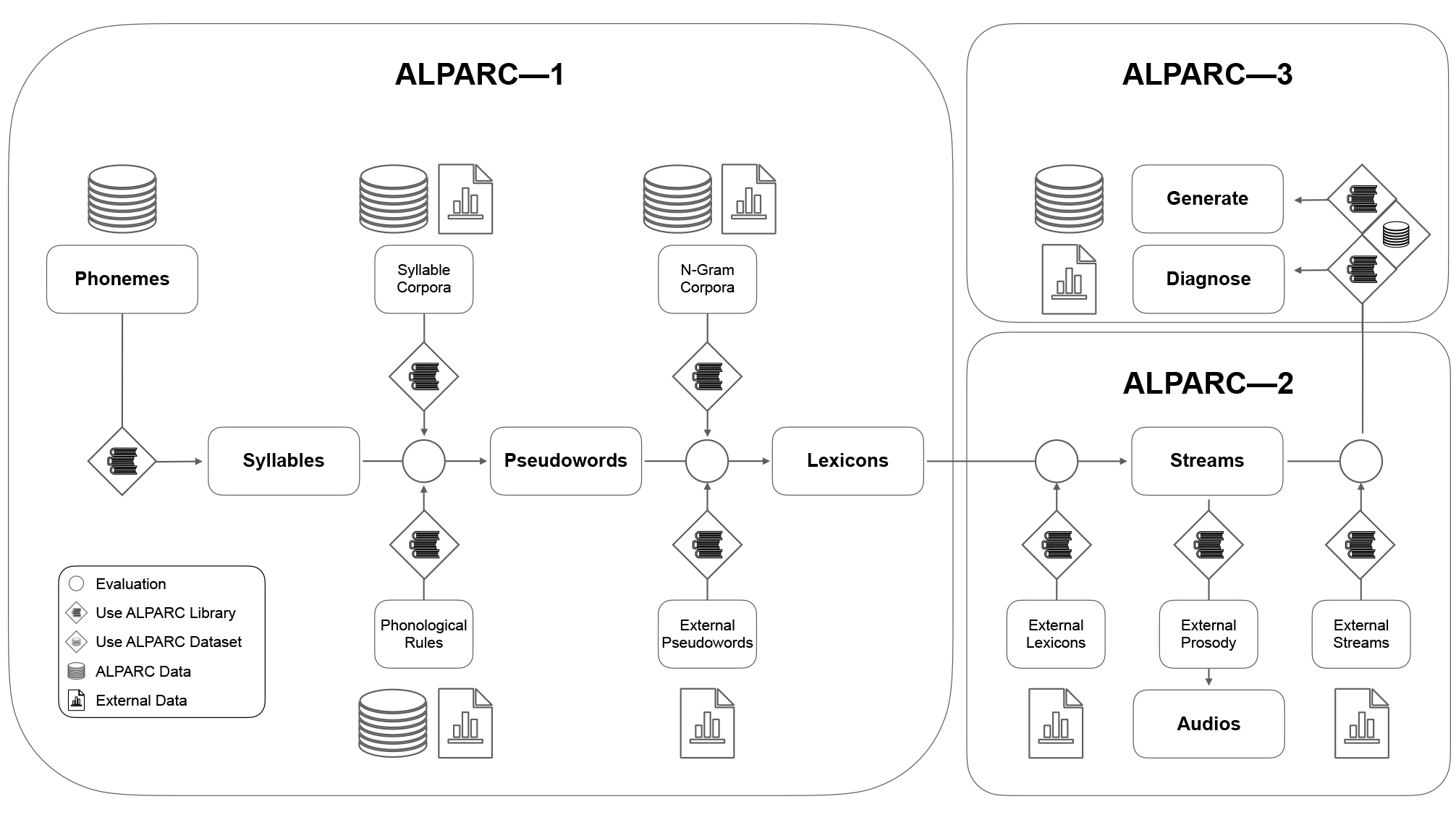
ALPARC workflow and modules. The pipeline highlights all the steps for the generation of artificial language streams. Phonemes annotated with binary vectors of phonological features constitute the atomic linguistic unit of ALPARC. Language-specific corpora can be used to filter the syllable and pseudoword register based on syllable frequency of use and phoneme n-gram probabilities in real languages. Pseudowords can be generated either internally (using our linguistic constraints) or externally. Then, ALPARC constrains the assembly of a Lexicon Register to minimize phonological feature regularities among pseudoword pairs that would confound TPs. Finally, lexicons are converted into TP-controlled streams of syllables. Beyond this linear workflow, ALPARC can be clustered in three modules ALPARC—1 covers the first three steps—from Phonemes to Lexicons. ALPARC—2 covers the step of stream generation with audio editing extensions. ALPARC—3 diagnoses the output of the previous modules in terms of phonological rhythmicity.

**Table 1.**
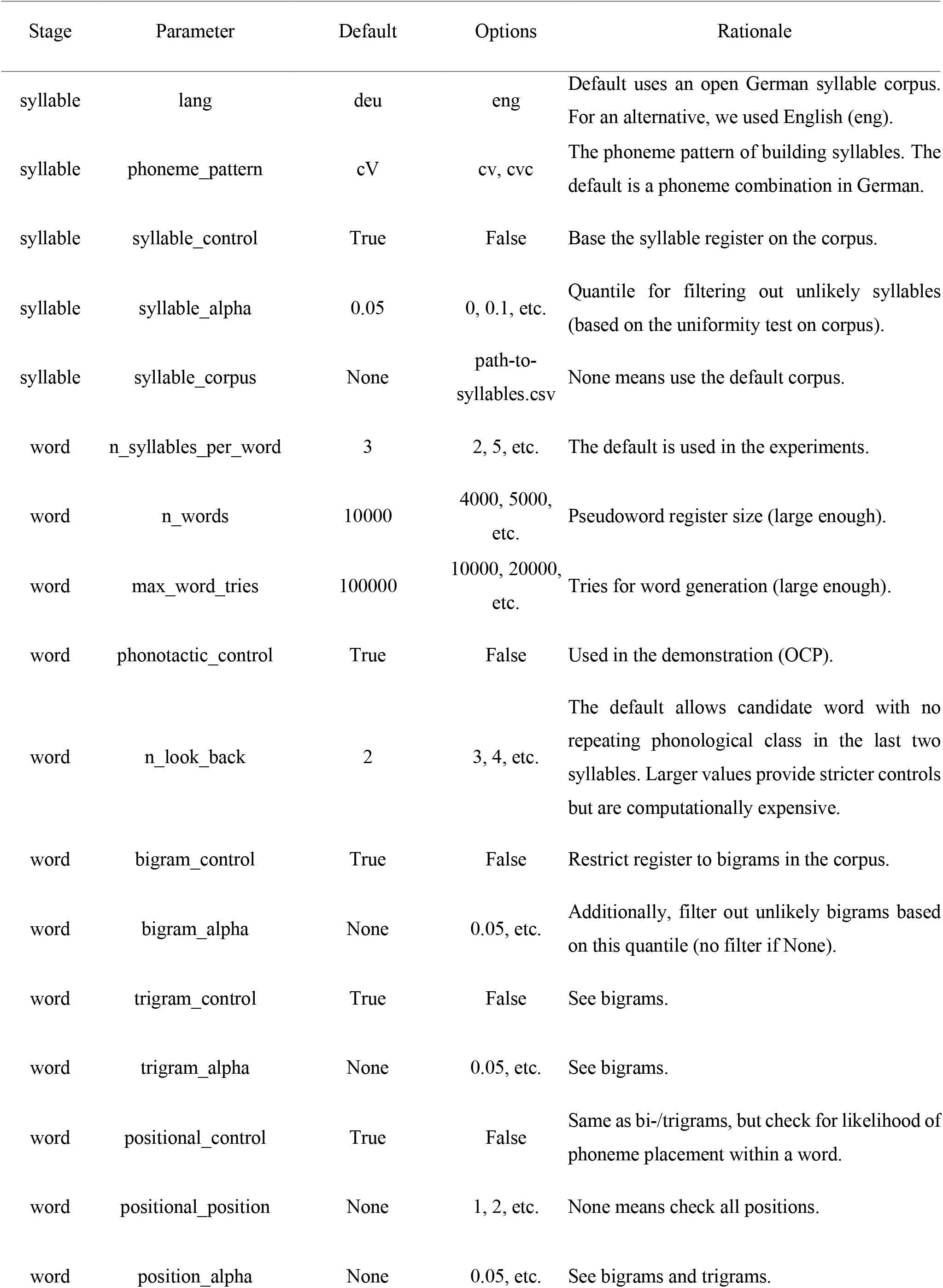
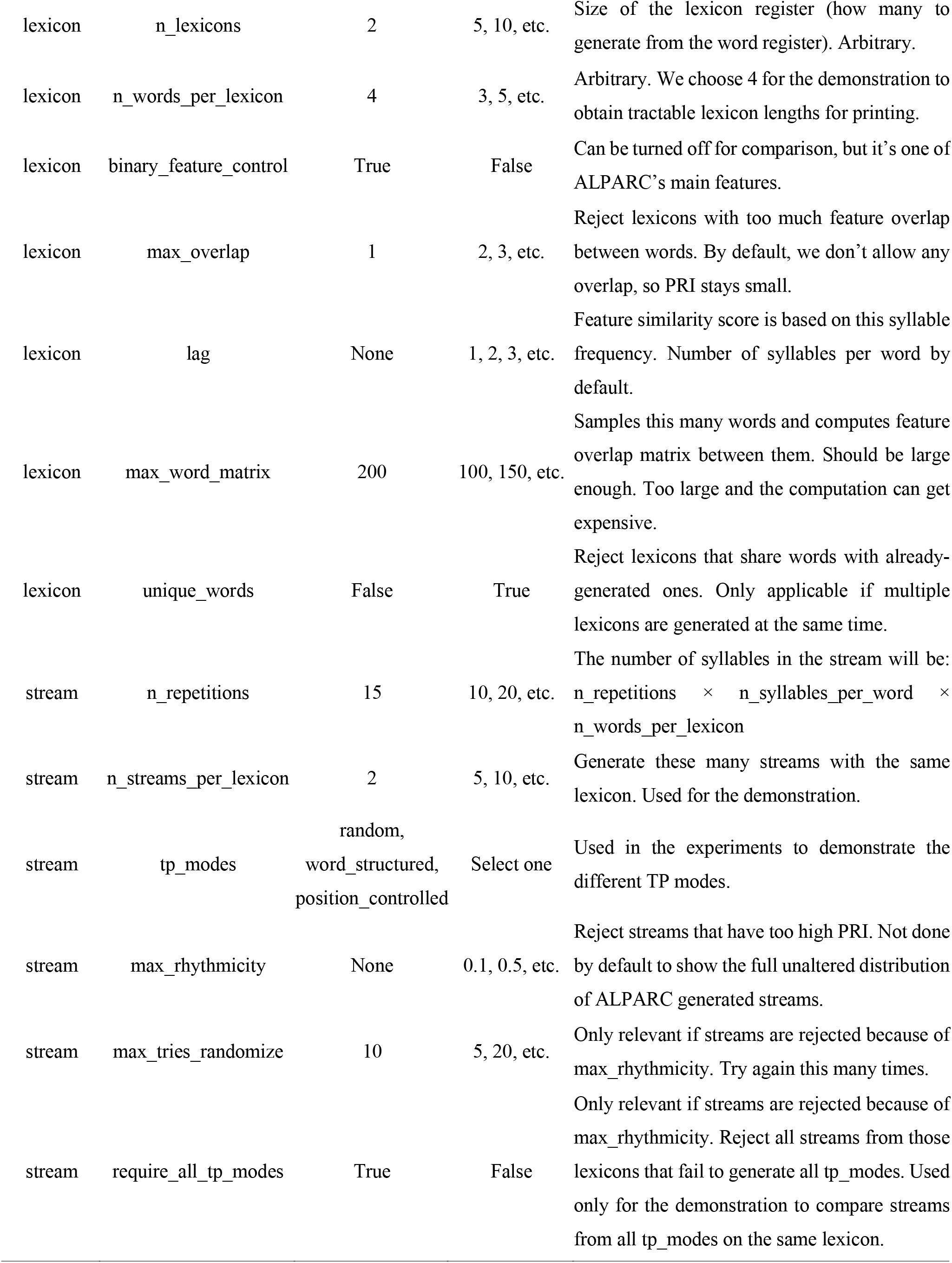
ALPARC parameter settings.

| Stage | Parameter | Default | Options | Rationale |
| --- | --- | --- | --- | --- |
| syllable | lang | deu | eng | Default uses an open German syllable corpus. For an alternative, we used English (eng). |
| syllable | phoneme_pattern | cV | cv, cvc | The phoneme pattern of building syllables. The default is a phoneme combination in German. |
| syllable | syllable_control | True | False | Base the syllable register on the corpus. |
| syllable | syllable_alpha | 0.05 | 0, 0.1, etc. | Quantile for filtering out unlikely syllables (based on the uniformity test on corpus). |
| syllable | syllable_corpus | None | path-to-syllables.csv | None means use the default corpus. |
| word | n_syllables_per_word | 3 | 2, 5, etc. | The default is used in the experiments. |
| word | n_words | 10000 | 4000, 5000, etc. | Pseudoword register size (large enough). |
| word | max_word_tries | 100000 | 10000, 20000, etc. | Tries for word generation (large enough). |
| word | phonotactic_control | True | False | Used in the demonstration (OCP). |
| word | n_look_back | 2 | 3, 4, etc. | The default allows candidate word with no repeating phonological class in the last two syllables. Larger values provide stricter controls but are computationally expensive. |
| word | bigram_control | True | False | Restrict register to bigrams in the corpus. |
| word | bigram_alpha | None | 0.05, etc. | Additionally, filter out unlikely bigrams based on this quantile (no filter if None). |
| word | trigram_control | True | False | See bigrams. |
| word | trigram_alpha | None | 0.05, etc. | See bigrams. |
| word | positional_control | True | False | Same as bi-/trigrams, but check for likelihood of phoneme placement within a word. |
| word | positional_position | None | 1, 2, etc. | None means check all positions. |
| word | position_alpha | None | 0.05, etc. | See bigrams and trigrams. |
| lexicon | n_lexicons | 2 | 5, 10, etc. | Size of the lexicon register (how many to generate from the word register). Arbitrary. |
| lexicon | n_words_per_lexicon | 4 | 3, 5, etc. | Arbitrary. We choose 4 for the demonstration to obtain tractable lexicon lengths for printing. |
| lexicon | binary_feature_control | True | False | Can be turned off for comparison, but it's one of ALPARC's main features. |
| lexicon | max_overlap | 1 | 2, 3, etc. | Reject lexicons with too much feature overlap between words. By default, we don't allow any overlap, so PRI stays small. |
| lexicon | lag | None | 1, 2, 3, etc. | Feature similarity score is based on this syllable frequency. Number of syllables per word by default. |
| lexicon | max_word_matrix | 200 | 100, 150, etc. | Samples this many words and computes feature overlap matrix between them. Should be large enough. Too large and the computation can get expensive. |
| lexicon | unique_words | False | True | Reject lexicons that share words with already-generated ones. Only applicable if multiple lexicons are generated at the same time. |
| stream | n_repetitions | 15 | 10, 20, etc. | The number of syllables in the stream will be:<br>$n\_repetitions \times n\_syllables\_per\_word \times n\_words\_per\_lexicon$ |
| stream | n_streams_per_lexicon | 2 | 5, 10, etc. | Generate these many streams with the same lexicon. Used for the demonstration. |
| stream | tp_modes | random,<br>word_structured,<br>position_controlled | Select one | Used in the experiments to demonstrate the different TP modes. |
| stream | max_rhythmicity | None | 0.1, 0.5, etc. | Reject streams that have too high PRI. Not done by default to show the full unaltered distribution of ALPARC generated streams. |
| stream | max_tries_randomize | 10 | 5, 20, etc. | Only relevant if streams are rejected because of max_rhythmicity. Try again this many times. |
| stream | require_all_tp_modes | True | False | Only relevant if streams are rejected because of max_rhythmicity. Reject all streams from those lexicons that fail to generate all tp_modes. Used only for the demonstration to compare streams from all tp_modes on the same lexicon. |

ALPARC provides a framework to model nested (linguistic) units for stimuli design in SL. Phonemes are the atomic linguistic units of ALPARC. These units are combined to form so-called registers of syllables, pseudowords and lexicons. We use the term register in its computational connotation referring to a constrained pool of candidates. ALPARC enables a fast search over large combinatorial spaces within the registers. The phonological domain of ALPARC is limited to 5935 phonemes with 21 phonological features each (Chomsky & Halle, 1968). These phonemes are identified in International Phonetic Alphabet (IPA) notation and further annotated with class features (syllabic [syl], sonorant [son], consonantal [cons]); manner features (continuant [cont], delayed release [delrel], lateral [lat], nasal [nas], strident [strid]); laryngeal features (voice [voi], spread glottis [sg], constricted glottis [cg]); and place features (i.e., anterior [ant], coronal [cor], distributed [distr], labial [lab], high [hi], low [lo], back [back], round [rnd], tense [tense], long [long]). Based on this matrix of binary phonological features linked to each phoneme, ALPARC generates pseudoword and lexicon registers of arbitrary size with one fixed syllable structure (e.g., [open] consonant-vowel syllables).

ALPARC can tailor an artificial lexicon to language-specific corpora distributions of syllable/phoneme (co-)occurrences. To that end, ALPARC requires a linguistic corpus that must be provided by the user. The tokenized entries in the corpus (e.g., phonemes or syllables) must be found in the phoneme list (Chomsky & Halle, 1968) and associated with their natural frequency of use, or phoneme statistics (e.g., bigram, trigram, or positional probabilities)., In the current application, ALPARC filters its lexical registers to approximate a more uniform distribution of the probability of (co-)occurrence of syllables and phonemes in spoken German (Arnold & Tomaschek, 2016; Schiel, 2010) and English (Baayen et al., 1996) corpora.

To summarize, ALPARC presents three modules that can be used independently and interdependently. ALPARC—1 generates ad-hoc artificial languages with linguistic and phonological controls. First, linguistic controls rest on syllabic and phonemic corpora, and special phonotactic principles (e.g., OCP). These features are employed to milden linguistic biases in artificial registers (Figure 2A-B). Second, phonological rhythmicity controls rest on the binary vectors of phonological features and return selected sets of pseudowords with a low feature overlap to reduce phonological confounds at the rate of TP boundaries (Figure 2C). ALPARC—2 makes three types of TP-uniform streams contrasting the information of each syllable at regular locations throughout the stream. To attain uniform TP distributions in a more precise and consistent way than with naïve shuffling implements acoustic and statistical rhythmicity controls solutions, ALPARC “walks” a Eulerian Circuit (EC). An EC is a closed walk on a graph that uses every directed edge exactly once. In our application, edges express local contingencies that correspond to TPs between syllables or pseudowords (i.e., nodes) in the artificial language (Figure 2D). To render such streams auditorily, we provide audio editing tools that concatenate syllables with a uniform duration, integrated with suprasegmental pitch samples (Figure 2E). ALPARC—3 expresses each stream of syllables as a binary matrix of phonological features across syllables. This representation allows to return a Phonological Rhythmicity Index (PRI). This index expresses the density of a (TP) pattern in each of the phonological feature vectors of a syllable stream (Figure 2C, right).

**Figure 2.**
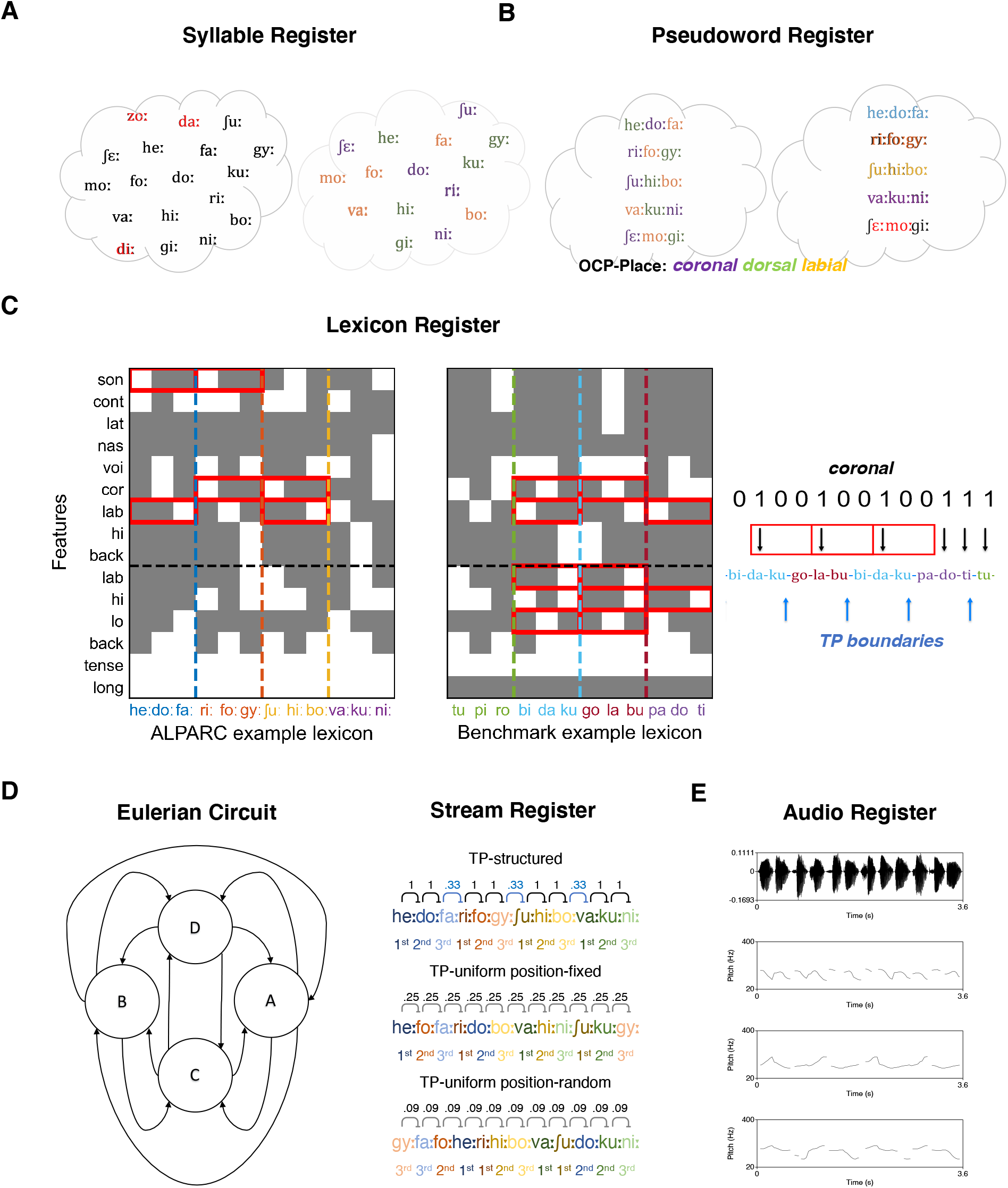
ALPARC registers. (A) Syllable register. Syllable tokens whose frequency deviate from uniformity in Schiel (2010) are excluded (red) from the register (left). The survival syllables are coded based on place of articulation (right). (B) Pseudoword register. One ad-hoc phonotactic constraint (i.e., OCP-place) shapes how syllables are allowed to form pseudowords (left). The resulting register is filtered (red) based on deviations from uniformity of phoneme bi-/tri-gram probabilities in Arnold & Tomaschek (2016). (C). Lexicon register. Binary matrices of phonological feature overlap across pseudowords pairs in one example lexicon generated with ALPARC (left), and one reference lexicon (right). Pseudowords are color-coded on the abscissae; vertical dashed lines cut the matrices at their boundaries. The horizontal line separates features associated with the consonant (top) or the vowel (below) phoneme of each syllable. Grey squares represent a minus (or a zero) in the feature vector of a given syllable (i.e., the feature is absent); whereas white squares represent a plus (or a one) for that feature (i.e., the feature is present). Red boxes flag repetitive patterns of phonological features across minimal pseudowords pairs that would confound the TP patterns at the pseudoword rate (e.g., [1 0 0, 1 0 0]). Phonological rhythms at the rate of TP boundaries (blue arrows) are to-be-avoided in the binary feature vector (right). To avoid the confound, ALPARC expands this feature matrix for all pseudoword pairs in the register to form lexicons with the least cumulative feature overlap. (D) Stream register. Eulerian circuit describing the TP uniformity (left) across four states (circles). Three uniform stream conditions can be formed with different patterns of TPs (right). TPs are displayed above neighboring syllable pairs. Pseudowords and syllable positions are color-coded in hue and shade, respectively. (E) Audio register. Sounds can be edited to have a fixed duration (top). In addition, the original pitch tier of the sound (mid-top) can be replaced, for instance, with a regular prosodic contour (e.g., bottom: adapted from Lamekina & Meyer (2023)).

## 3. Method

The prime use-case of ALPARC is to build artificial languages of pseudowords step-by-step. First, ALPARC connects phonemes into syllables (Step 1) and pseudowords (Step 2), while constraining legal compositions linguistically through ad-hoc phonotactic rules and speech corpora. Next, ALPARC builds lexicons as sets of pseudoword candidates with minimal phonological regularities (Step 3). Lexicons are rendered as sequences of tokens with uniform TP distributions across all syllables or across regular pseudoword boundaries (Step 4). Audio editing tools optimized for frequency-tagged streams are available (Step 5).

### 3.1. Step 1—from phonemes to syllables: lexical control

In Step 1, ALPARC builds a phoneme register, annotated with binary phonological feature vectors. By default, ALPARC selects a subset of phonemes from a universal corpus (Chomsky & Halle, 1968). ALPARC can thus build a phoneme register that is limited to the phonemes in the list, and may not cover all of the speech sounds of all of the world’s languages. In our implementation, we restrict the phoneme register to retain only single IPA characters with no coarticulation. This design choice was meant to ease downstream computations, albeit more complex (e.g., coarticulated) phonemes are present in the matrix and can be selected.

To compose the syllable register, ALPARC uses one fixed syllable structure. The default (cV syllables, with long vowels) is set to render vowel duration uniform across syllables, for isochronous manipulations (e.g., frequency-tagging). While selecting the syllable type, users may also specify a subset of phonological features for each phoneme (default = sonorant [son], continuant [cont], lateral [lat], nasal [nas], voice [voi], coronal [cor], labial [lab], high [hi], back [back] for consonants [C]; and labial [lab], high [hi], low [low], back [back], tense [tense], long [long] for vowels [V]). These features will be carried over as annotations for further steps.

ALPARC can filter a syllable register to approximate a uniform distribution from linguistic corpus data. In particular, syllables whose frequency of use significantly deviate from the uniform distribution are removed (Figure 2A). Corpora-based filters can be useful for two reasons. First, using real syllables from the native language of the learners is essential to ensure intelligibility, since non-distinctive phonemes are ambiguous. Second, eliminating highly (in)frequent syllables would reduce the potential tendency to endorse an artificial pseudoword due to the presence of a particularly (un)familiar syllable. Users can also skip this step and import an externally-generated pseudoword list to have their target pseudowords returned as an annotated register.

### 3.2. Step 2—from syllable to pseudowords: phonotactic control

In Step 2, ALPARC generates a pseudoword register from a syllable register with a fixed number of syllables (default = 3). In line with the previous step, the pseudoword register may be filtered based on whether phoneme n-grams are present in the corpus (default = True), and the threshold to approximate uniformity can be adjusted (default α threshold = none [i.e., without excluding the tokens that deviate from uniformity]). This filter is set to ensure that the resulting register contains tokens that belong to a target language. Moreover, the pseudoword register may be filtered based on positional statistics (positional_control = True), removing pseudowords that present a phoneme that rarely occupies a certain word-position (positional_position = None [i.e., all]).

In addition to these filters, ALPARC can apply phonotactic constraints within and across pseudowords. To that end, ALPARC combines simple phonological features into phono-articulatory classes (Ramers, 1998), that are divided into balanced categories according to their place of articulation (labial, apical, other), manner of articulation (sonorant, plosive, fricative), and vowel identity. In particular, users may set the degree of phonotactic control to OCP-place (Boll-Avetisyan & Kager, 2014), -manner, and vowel identity constraints. OCP constraints on these articulatory features are then used to restrict the syllable combinations that are allowed to form pseudowords (Figure 2B). By default, ALPARC generates trisyllabic pseudowords according to all three of these rules, meaning that, the resulting pseudowords will comprise three syllables with a different place and manner of articulation of the consonants and different vowels. This feature has been designed for German, but it can be disabled or adjusted (n_look_back) if the input register is too small. Alternatively, users can add custom code blocks to control for different phonotactic rules based on their target language.

ALPARC interprets user input to return annotated lexicons and streams. For instance, users may input pseudowords generated with external tools, such as Wuggy (Keuleers & Brysbaert, 2010), which might be suitable for language-specific phonotactic controls. To enter our pipeline, externally-generated registers must be akin to those that ALPARC would generate by itself. Thus, all pseudowords should have a fixed syllable structure and a fixed number of syllables/phonemes. An interpretable input may comprise a set of pseudowords as split (“|”) strings of syllables, in IPA, with special phoneme separators (“_”). As long as all input phonemes are present in our list, (a subset of) their phonological features can be retrieved for further rhythmicity controls.

### 3.3. Step 3—from pseudowords to lexicons: phonological controls

In Step 3, ALPARC generates lexicon registers preventing phonological rhythmicity. Users may specify the number of pseudowords per lexicon (default = 4), the number of lexicons to be generated (default = 5), and if such lexicons should be comprised of all unique words (default = False). In addition, ALPARC uses a subset of pseudowords (default = 200) in order to speed up further computations. The output lexicon registers contain sets of pseudowords with minimal feature overlap to reduce phonological regularities at the pseudoword rate.

Repetitions of the same phonological features at regular positions within a syllable stream may randomly accompany TPs and confound SL. In other words, if neighboring pseudowords share a phonological feature at the same phonemic position, but not at all other positions, that feature will be rhythmic and confound the TP rate. To tackle this issue, we reasoned that limiting the opportunity for a feature to occur at regular intervals will reduce its periodicity in structured streams. Since pseudowords occur at a fixed fraction of the syllable rate (e.g., 1/3 for trisyllabic tokens), ALPARC builds a matrix of position-wise phonological of feature overlap across pseudoword pairs. For each pseudoword pair and each phonological feature, shared binary sequences that mimic the regular TP pattern are flagged (Figure 2C). ALPARC avoids rhythmic patterns among syllables that occupy the same relative position between neighboring pseudowords.

ALPARC uses a lazy evaluation strategy to prevent phonological rhythmicity at the lexicon level. At that level, lazy evaluation means that pseudoword combinations with the least feature overlap are generated first, so that lexicons are only created when they are referenced in subsequent steps. Next, ALPARC derives a square matrix of feature overlap as the sum of flags across features for each pseudoword pair. Based on this matrix, ALPARC ensures minimal cumulative feature overlap. This strategy minimizes computation time relative to generating all combinations of pseudowords.

It is here important to mention once again that ALPARC can read phonemic transcripts. At this stage, users may input a phonemic transcript (strictly in X-SAMPA or IPA format) based on their own artificial or natural speech recordings and retrieve a subset of phonological features for each phoneme. ALPARC controls phonological rhythmicity at a fixed rate by having full control over the matrix of phonemic features, such that they can be combined arbitrarily. For instance, if coarticulated phonemes are included, users might code ad-hoc rules for how phonemes are allowed to combine, similarly to how we did with the OCP. As a caveat, nominal consistency of the phonological feature set for each syllabic must be kept for phonological rhythmicity controls. For instance, users may add a nominal amount of extra binary feature vectors (e.g., expressing a given phonological class, such as fricative, or a type of coarticulation, hesitation, or stress. These additional arbitrary features would need to be added manually to the binary feature matrix as row vectors (by adding a [1] when present, in a vector of [0]s).

### 3.4. Step 4—from tokens to chains: TP control

In Step 4, ALPARC creates sequences of tokens with uniform TP distributions. To that end, we considered a random shuffling of a fixed number of tokens to be exposed to some transitions being more likely than others due to randomness. As a result, TPs would be biased because tokens present different cue diagnosticity on the microscale of all individual pairs. Furthermore, under loose constraints, TPs may vary throughout the stream, leading to irregularities at the macroscale of the stream, with different TPs for the same pair at different stream locations. Statistical irregularities at the micro and macroscales are not exempt from having an impact on neural responses (Hasson, 2017). If TPs are variable or time-varying, certain boundaries can be perceived as more or less salient than others, with behavioral and neural consequences in terms of learning and tracking. To minimize this statistical confound, ALPARC uses a different method that guarantees TP stationarity.

ALPARC generates a finite sequence *V* = (𝒱_1_, 𝒱_2_, …, 𝒱_r_) from an alphabet of size *N* such that: (i) every syllable appears equally often, (ii) no syllable immediately repeats, and (iii) every ordered pair of distinct syllables (i → j) occurs at the uniform target count:

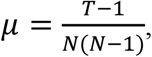

meaning that the empirical transition probability from any syllable i to any syllable j ≠ i is as close to 1/(N−1) as a finite sequence allows. Any sequence satisfying Conditions (i) – (iii) is equivalent to a Eulerian Circuit (EC) that uses every directed edge exactly once. We construct the graph in a particular way where vertices are syllables, and edges (i → j) are added with multiplicity μ, distributed so that every vertex has equal in-degree and out-degree (Figure 2D, left). This is technically a directed multi-graph with balanced edge counts. Reading off the vertices of the EC enables ALPARC to produce the syllable stream with transition counts equal to the edge multiplicities by construction. Using Hierholzer’s algorithm (Corberán & Laporte, 2015; Hierholzer & Wiener, 1873), ALPARC follows edges until stuck, then splices in detours at vertices with unused edges, until all edges are consumed. A greedy minimum-count rule improves the speed of convergence to the desired TPs: Always pick the outgoing edge (i → j) whose current count is smallest, breaking ties at random. This problem is similar to drawing all solutions of the “Haus vom Nikolaus” (Hoch & Küveler, 2023) consecutively without repeating any solution twice before all others.

With this logic, ALPARC makes TP uniform streams of symbolic tokens, reflecting pseudowords. (Figure 2D). Tokens repeat a number of times (default = 15). To ensure TPs remain constant throughout the stream, ALPARC samples new tokens iteratively based on a memory matrix of TPs. More specifically, the sampling of new tokens is limited among those that have the lowest TP to the previous token based on the memory matrix. As a result, syllable-tokens in position-fixed streams are constrained to always “walk” the viable path they have walked the least, based on the memory matrix. Moreover, every token must be used once before any could be repeated more than all others and consecutive repetitions of the same token are not allowed. Thus, ALPARC produces streams with stable TPs across boundaries. When ALPARC expands pseudowords into plain sequences of syllables, TPs between syllables within any pseudoword are always 1 (Figure 2D, top).

ALPARC adopts a similar approach to attain uniform TPs across all adjacent syllables. In fact, ALPARC has two different functions for the TP-uniform position-random and the TP-uniform position-fixed streams. In these conditions, each token maps directly to a syllable. In position-fixed baseline streams syllable-tokens are constrained to occupy only one fixed position within a pseudoword-token, and to follow one another respecting this positional rule, where all syllable TPs tend to exactly 1⁄*N* throughout the stream (Figure 2D, center). In TP-uniform position-random streams, syllable TPs are equal across positions within a pseudoword-size chunk with no consecutive repetitions allowed, leading to TPs of 1⁄*N* − 1 (Figure 2D, bottom).

Surprisal, entropy, and non-uniformity metrics quantify information in the three stream modes. At time t, the empirical distribution over next syllables 𝒱_t+1_ = *j* given the current one 𝒱_t_ = *i* is the empirical average of transition counts observed before *t*:

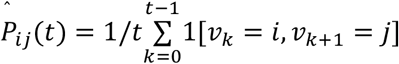

The surprisal of observing the actual next syllable 𝒱_t+1_ is:

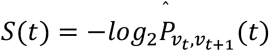

So, the conditional entropy of the transition distribution from source I (with convention log 0 = 0) is:

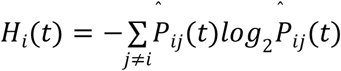

Averaging over sources yields *H* (*t*), whose maximum is log_2_(N−1) (uniform transitions). Similar to entropy, the KL-Divergence measures the distance to the uniform distribution:

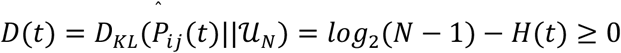

This non-uniformity metric collapses to zero exactly when empirical TPs are perfectly uniform.

ALPARC makes three types of stream variants with different local TP distributions. Across variants, ALPARC ensures that each token follows every other token equally often to precisely manipulate TPs between pseudowords in TP-structured streams and TPs between syllables in TP-uniform streams. The three types of streams that ALPARC generates contrast the information at word boundaries. First, the TP-structured variant is built at the word level, where each word is a single vertex and later substituted as syllable strings. Here, TPs across word boundaries are uniform (1/(W−1)), whereas TPs are 1 within words, providing the segmentation cue. Second, the TP-uniform position-random variant is constructed as a complete directed graph on all N syllables; where each edge (I → j), I ≠ j, gets multiplicity ⌊μ⌋ or ⌈μ⌉. Here, any syllable can follow any other with equal probability 1/(N−1). Third, the TP-uniform position-controlled variant is built such that syllables are partitioned into position groups, where edges only point from and to adjacent groups, and each cross-group pair gets exactly R edges. Here, TPs are uniform within each position transition, and all W targets are equally likely (1/W), while syllables only appear at their designated within-word position.

### 3.5. Step 5—from languages to sounds: rhythmicity control

In Step 5, ALPARC uses ad-hoc audio editing tools to produce frequency-tagged auditory streams. We first synthesized individual syllables into separate audio files using Wavenet (Oord et al., 2016), and the Speech-Synthesis-Markup-Language (Taylor & Isard, 1997). Praat (Boersma & Weenink, 2007) was used to edit the audio files of individual syllables. Here, we equalize syllable duration to 300 ms and their amplitude to 65 dB. Furthermore, ALPARC affords pitch contour manipulations for frequency-tagging stimuli. Naturalistic pitch contours can be separately recorded or synthesized (see also Lamekina & Meyer, 2023) and edited to an arbitrary duration that matches the size of syllables or pseudowords. Thence, suprasegmental cues can be added to the original syllable stream by replacing its pitch tier with a new prosodic contour (Figure 2E).

ALPARC assesses confounding patterns at the TP rate, using a Phonological Rhythmicity Index (PRI). To demonstrate how the PRI is calculated, let us assume a stream of four trisyllabic pseudowords. In this stream, TP patterns repeat periodically every three syllables, with TPs of 1 within pseudowords and .33 at pseudoword boundaries. Such a sequence can be recoded into a binary vector of “TP decays” marking low-TP events as 1 and high-TP events as 0 (i.e., [0 0 1 0 0 1, …]). Based on this TP pattern, we define kernels as vectors that represent two instances of TP decays at all possible lags (i.e., [1 0 0 1 0 0], [0 1 0 0 1 0], [0 0 1 0 0 1]). This definition rests on the assumption that a sequence must repeat consecutively at least twice, for it to be classified as a pattern. Now, let us assume one vector *k* = [1 0 0 1 0 1 0 0 1 0 0 1 1 0 0], representing the presence (1) or absence (0) of one feature throughout a stream. ALPARC convolves the three kernels through the feature vector *k*, assessing whether any of the kernels matches the current portion of the feature vector. For instance, at the first iteration—*k* _*(1,6)*_ = [1 0 0 1 0 1]—there is no exact match with any of the kernels; hence, ALPARC notes a non-match as a zero. One step ahead, *k* _*(2,7)*_ = [0 0 1 0 1 0], there is again no match. The fifth iteration, *k* _*(5,10)*_ = [0 1 0 0 1 0], presents an exact match to one of the kernels; so, ALPARC notes a one. By the end of the stream—until *k* _*(10,15)*_—ALPARC reports a total of 3 matches over 10 iterations. The PRI is the ratio between the sum of all such matches and the total number of comparisons. In this example, the value of .33, indicates that 33% of the pseudoword tracking effect that might be attributed to TP patterns may be due to phonological regularities of the feature under examination. A lower PRI denotes a lower density of rhythmic phonological patterns at the TP rate.

In sum, the PRI measures the average periodicity of all phonological features at the TP rate of interest in the syllable streams. The PRI is computed on each feature vector by convolving the binary TP kernels. Effectively, the kernels were designed to represent two instances of a TP pattern (e.g., [1 0 0 1 0 0]) at the rate of interest. ALPARC uses the PRI as a descriptive metric of confounding phonological regularities at the rate of rhythmic (TP) kernels. The maximum PRI across features assesses the worst-case scenario of phonological regularities in a stream. In contrast, autocorrelations cannot resolve the presence of exact binary patterns.

## 4. Results

Our results suggest that ALPARC lexicons reduce linguistic, statistical, acoustic, and phonological confounds. Before showing these results, the next section covers how we evaluated the output of ALPARC at each stage. All our findings resulting from the evaluation of each step are detailed in separate sections.

### 4.1. Evaluation

First, we demonstrate how ALPARC filters a register based on syllable frequency of use and phoneme n-gram probabilities extracted from linguistic corpora. The frequency or probability distribution associated with the tokens in the corpus is log-transformed, normalized (z-score), and tested against the uniform distribution. The p-value distributions stemming from the uniformity test are used to filter the syllable or the pseudoword register. This filter allows for normalizing the frequency distributions of syllables and phoneme n-grams in our syllable and pseudoword registers, meaning that the survival pseudowords do not contain too (in)frequent phoneme bigrams or trigrams.

To assess TP precision at the stream level, we used non-uniformity, surprisal, and entropy metrics. We plot these information metrics as a function of stream location for conditions. We compare our approach to a naive pseudo-randomization that entails shuffling of the tokens with the only constraint that consecutive repetitions of the same token are not allowed. We used Levene’s tests comparing the two strategies in terms of TP variance at pseudoword or syllable boundaries in TP-structured and TP-uniform streams, respectively. In addition, we synthesized the three types of streams and computed their amplitude modulation spectra using MATLAB (The MathWorks, Inc.).

To evaluate phonological controls, we compared different types of artificial languages of pseudowords. We generated 21 lexicons of pseudowords based on German corpora (—DEU), 21 lexicons based on English corpora (—ENG), and 21 random lexicons with no controls (—RND). We compared the PRI of these lexicons against 21 artificial lexicons (BENCHMARK) from the prior literature (Assaneo et al., 2019; Batterink & Paller, 2017; Cunillera et al., 2006; Kiai & Melloni, 2021; McNealy et al., 2006; Saffran et al., 1996). All these lexicons are made of four tri-syllabic pseudowords (Table S1). These criteria ensure a fair comparison with the lexicons that ALPARC produces by default.

To evaluate whether ALPARC reduces phonological rhythmicity, we compared different the PRI across lexicon types. For each lexicon (n = 21 per type), we simulated 10 streams per TP mode. We compared the PRIs across conditions using two-sided Welch’s independent sample t-tests. We used this non-parametric test since data arise from mixture distributions with non-Gaussian components. Here, the components are the distributions of stream-level PRIs generated from a single lexicon. While the resulting mixture is again non-Gaussian, Welch’s t-test remains valid for comparing means. This test does not assume normality of the data, but only that the sampling distribution of the mean is approximately normal, which is guaranteed by the Central Limit Theorem for our sample size of 21 lexicons with 10 streams each = 210 data points (Lumley et al., 2002). This asymptotic justification holds regardless of whether the population distribution is a mixture. We applied a Bonferroni correction of the p-values for the tests (Table 3).

For validation, we performed control analyses with different parameter settings. More specifically, we built streams in all TP modes with different number of syllables (2, 3, or 4) or pseudowords (3, 4, or 5) to show their PRI distributions (Table S2). To evaluate whether ALPARC feature controls prevent a high PRI, we computed the Pearson’s correlations between the feature overlap of ALPARC lexicons and the maximum PRI in the streams.

### 4.2 ALPARC registers adapt to linguistic corpora

Linguistic confounds that emerge from prior linguistic background are mitigated by approximating a uniform distribution from Zipf-distributed (Zipf, 1932) corpora statistics. The original range of frequency of use of the syllables is reduced from 1–19303 to 9–642 in German syllables in Schiel (2010) and from 1–72971 to 3–94 in English syllables in Baayen et al., (1996)’s corpus (Figure 3). Likewise, we show how ALPARC could filter a pseudoword register using bigram, trigram, and positional phoneme statistics based on a German corpus (Arnold & Tomaschek, 2016). By applying these filters, we produced two syllable and pseudoword registers— ALPARC—DEU and ALPARC—ENG—from which we sampled pseudowords with phonological controls.

**Figure 3.**
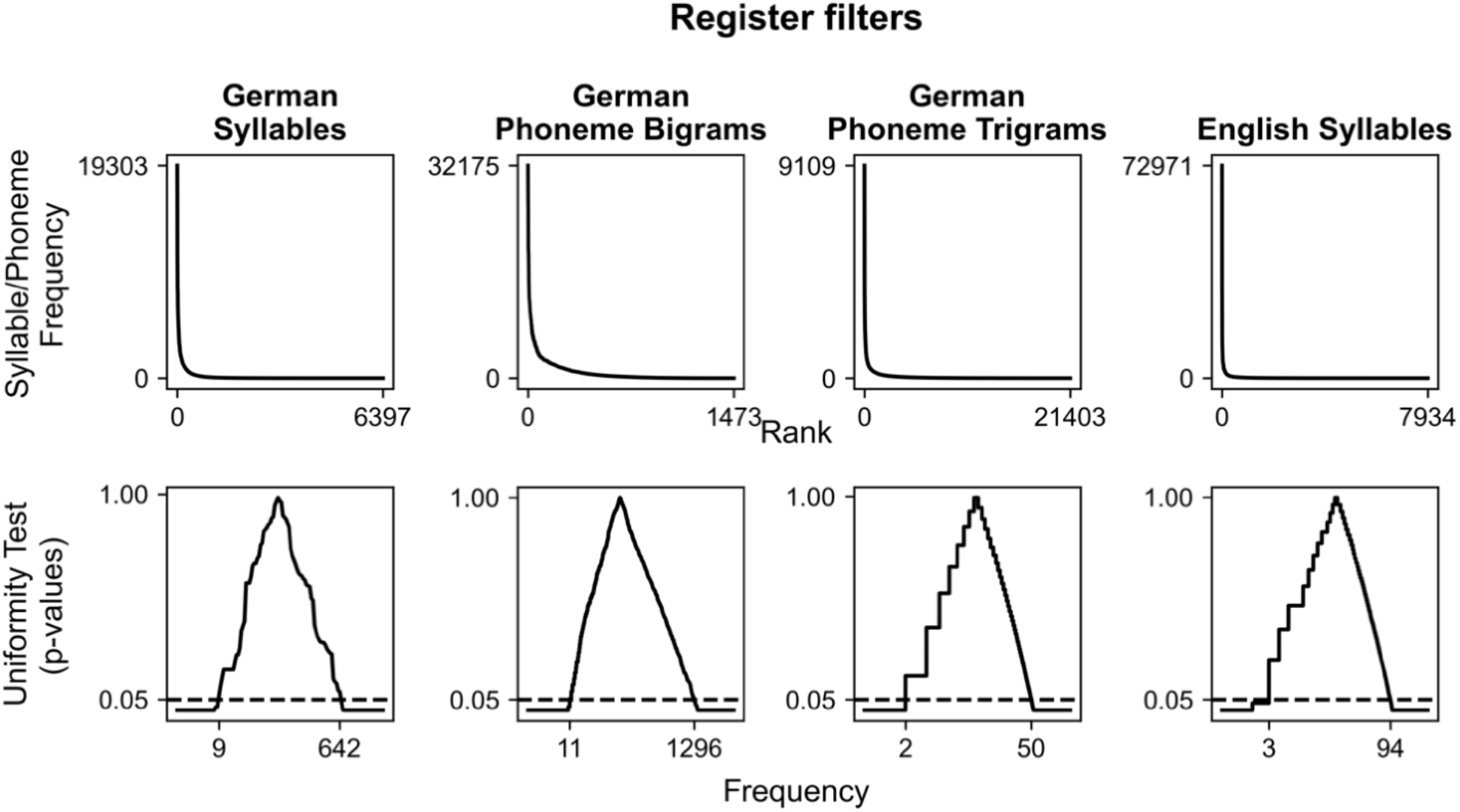
Register filters. Corpora-based distribution of syllable frequency of use or phoneme n-gram probabilities are plotted as a function of frequency-ranked tokens (top), above the distribution of p-values derived from a uniformity test based on the z-transformed frequency distribution above as a function of token frequency (bottom). The horizontal line represents an arbitrary alpha threshold (.05). German syllable distributions are based on Schiel (2010), while phoneme n-gram distributions are based on Arnold & Tomaschek (2016). English syllables are based on (Baayen et al., 1996).

### 4.3. ALPARC generates uniform TP sequences with acoustic controls

ALPARC generates syllable streams with uniform TP distributions. ALPARC attains TP uniformity relative to a random shuffling method (Figure 4). Levene’s tests show that ALPARC reduces the overall TP variance for both the TP-structured (*W* _(1, 476)_ = 31.72, *p* < .001), and the TP-uniform position-random streams (*W* _(1, 1436)_ = 142.76, *p* < .001). These results suggest that all types of streams generated with ALPARC reduce TP variance compared to randomly shuffled streams within each stream.

**Figure 4.**
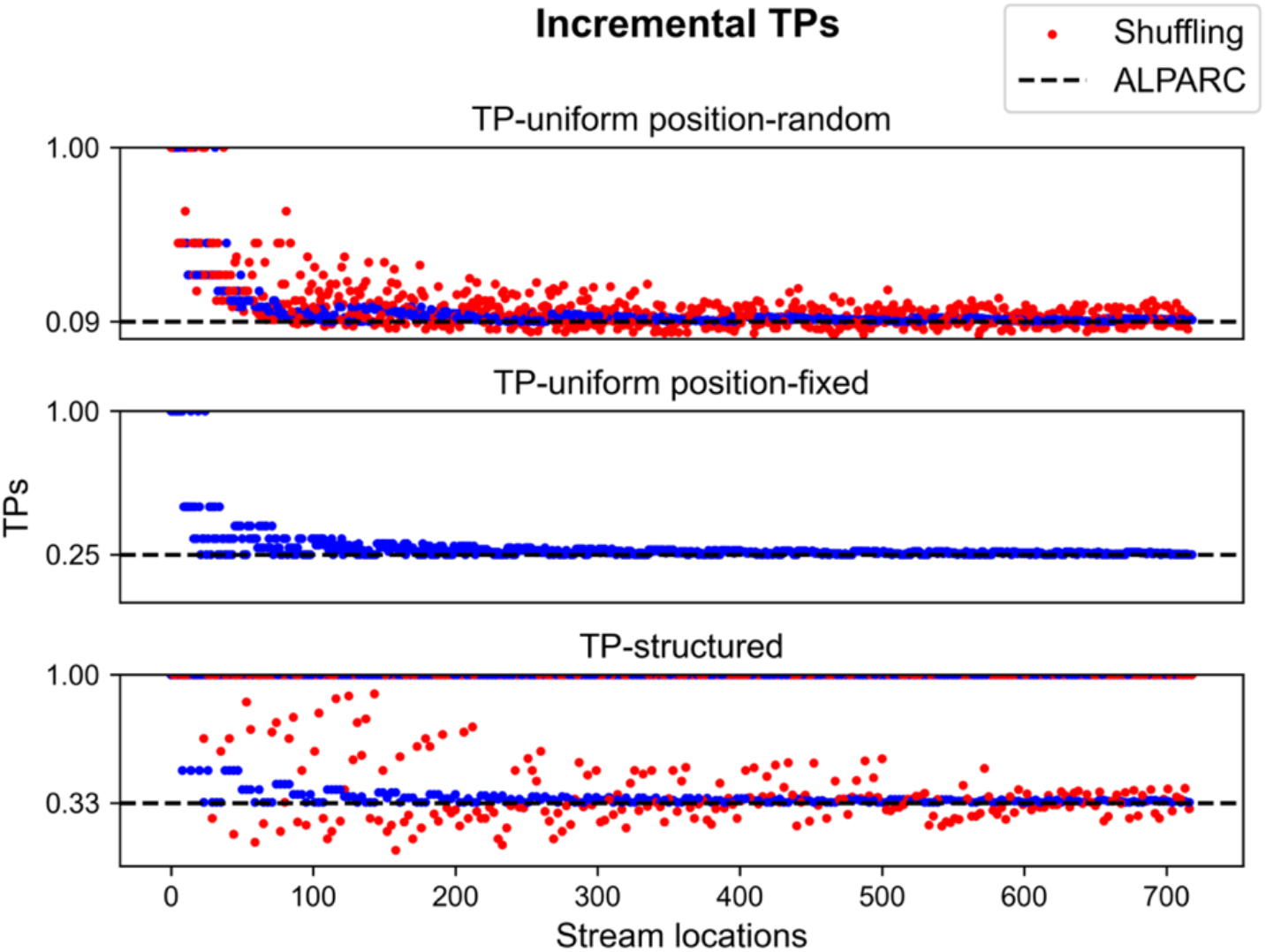
TP variance. Scatter plots of the observed TP values (colored dots) at each stream location, based on different strategies (red: Shuffling; blue: ALPARC). Dotted lines represent the theoretical value of TPs for a trisyllabic lexicon of four pseudowords.

The three stream modalities impose fundamentally different surprisal values at word boundaries (Table 2). Early in the stream, surprisal is volatile, but our greedy minimum-count algorithm converges quickly around the target value log_2_(N−1), in TP-uniform position-random streams. In TP-structured streams, within-word transitions have true probability 1. Therefore, surprisal under a structure-aware model is 0 mid-word and jumps to log_2_(W−1) at every word boundary. To corroborate this result, we calculated the empirical surprisal values across stream locations (Figure S1).

**Table 2.** TP stream modalities and associated information metrics.

| TP mode | Within-word surprisal | At word boundary | TP boundary? |
| --- | --- | --- | --- |
| TP-structured | 0 (true TP = 1) | $\log_2(W-1)$ | Yes. Maximum Contrast |
| TP-uniform position-random | $\log_2(N-1)$ | $\log_2(N-1)$ | No |
| TP-uniform position-fixed | $\log_2(W)$ | $\log_2(W)$ | Position identity only |

**Table 3.**
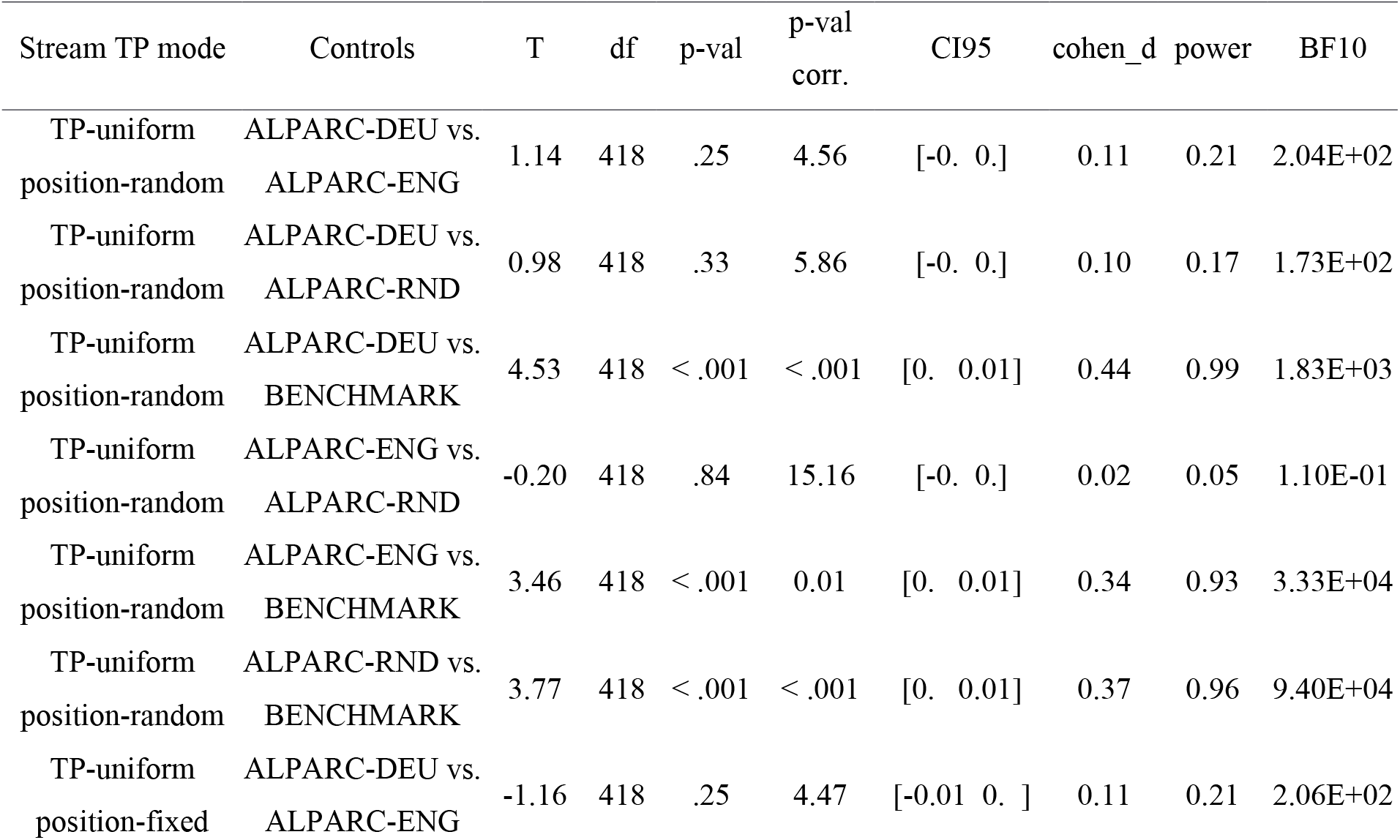
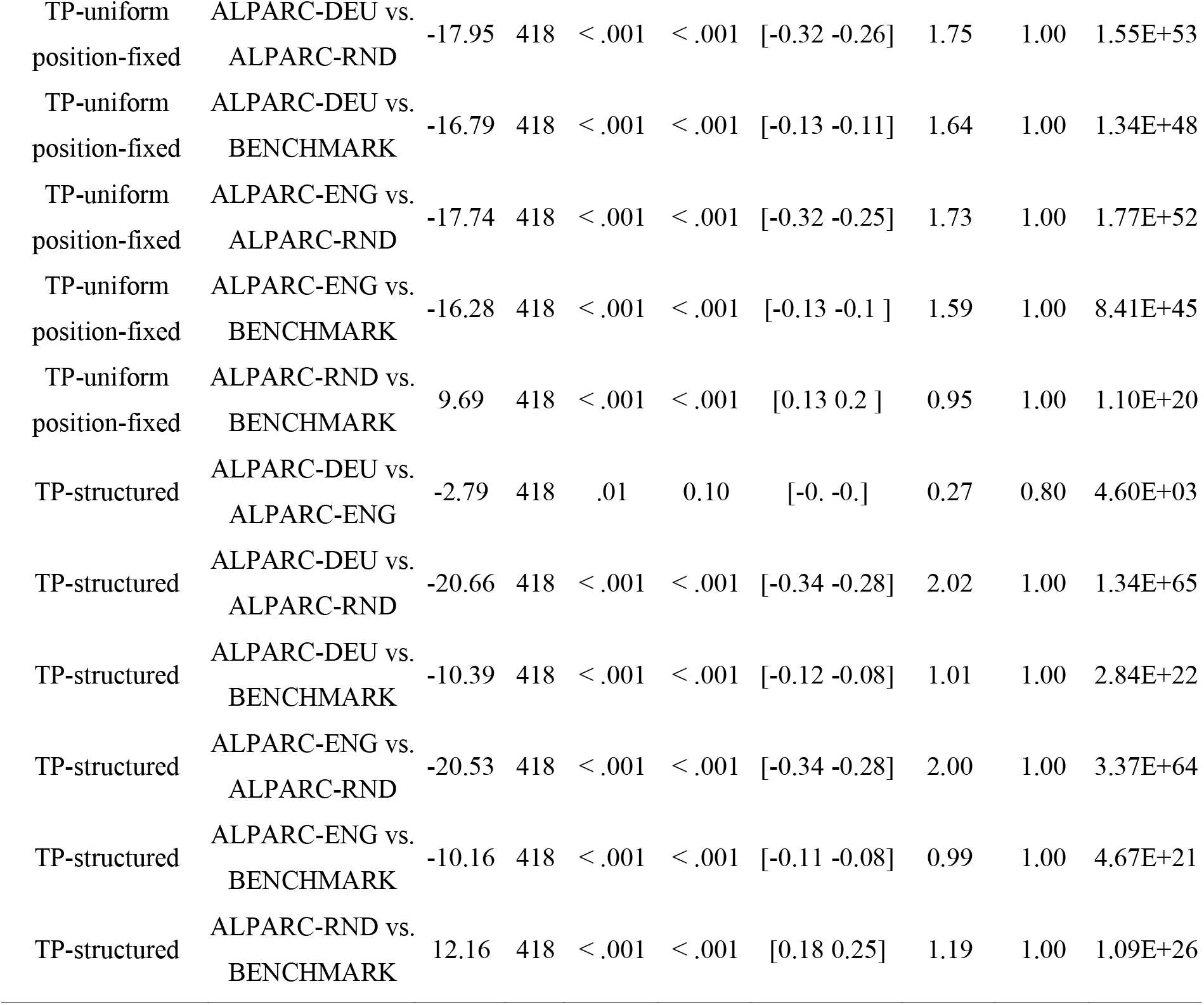
Independent Pairwise Welch’s t-test results of maximum PRIs for all stream TP modes.

In parallel, ALPARC factors out acoustic rhythmic confounds in two stream TP modes. To show which types of TP streams exhibit different spectral peaks at the pseudoword rate, we synthesized frequency-tagged auditory streams of syllables in each TP mode based on one example ALPARC lexicon (Figure 5). The amplitude modulation spectrum of the TP-uniform position-random stream displays no peak at the pseudoword rate (1.11 Hz); while the TP-structured and TP-uniform position-fixed spectra show the same peak at that rate.

**Figure 5.**
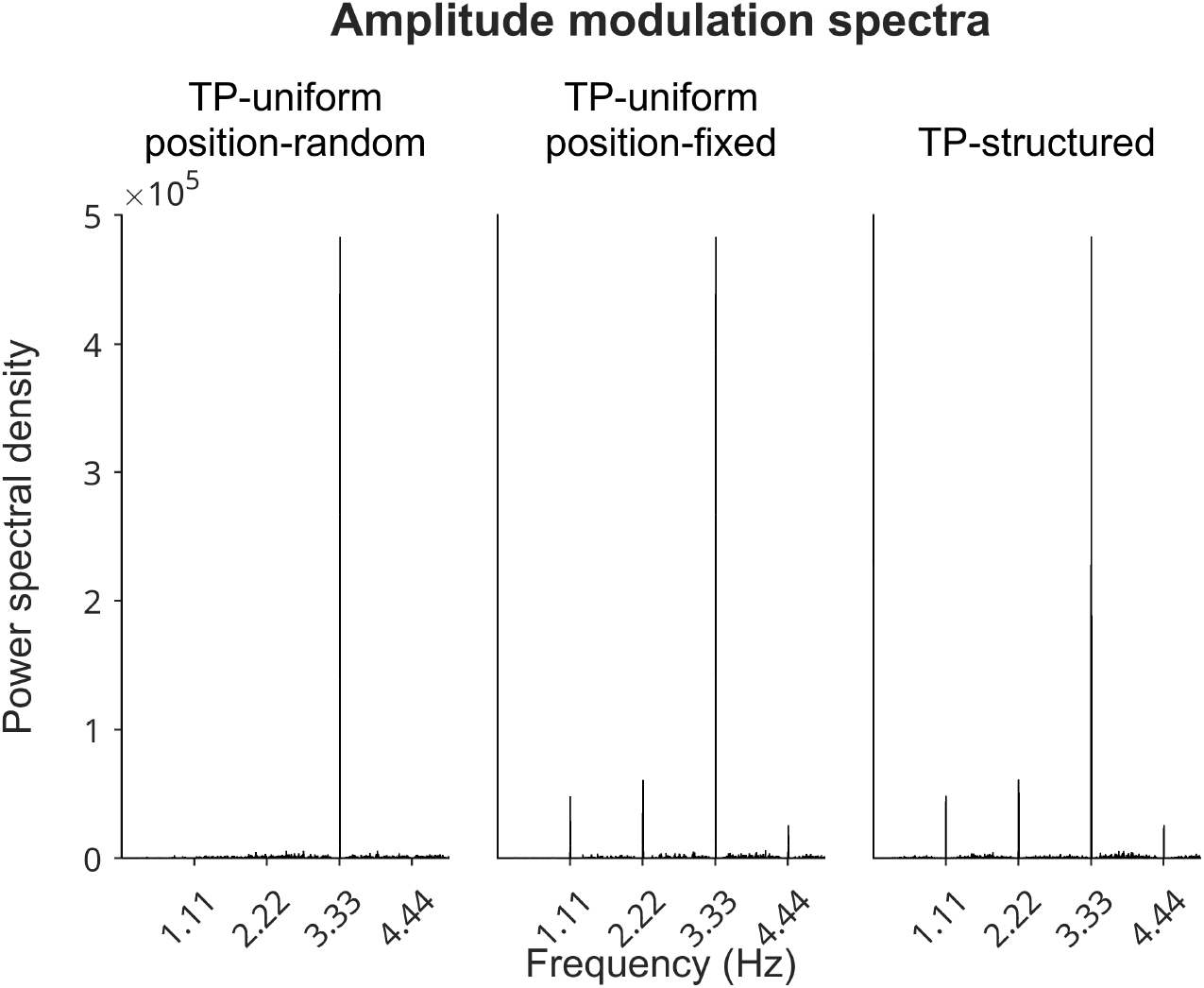
Speech modulation spectra. Power spectra from one example ALPARC lexicon in different stream TP modes. Each syllable had a fixed duration of 300 ms, leading to a peak at the syllable rate (3.33 Hz) in all modes. Syllables in the position-random stream occupy any possible location in the stream, leading to no peak at the rate of pseudowords (1.11 Hz). Syllables in the position-fixed and TP-structured streams are enforced to occur at fixed locations that are multiples of three, leading to a peak at the pseudoword rate (1.11 Hz), and its harmonic (2.22 Hz).

### 4.4. ALPARC lexicons reduce the PRI at the TP rate

ALPARC yields a low and robust PRI across streams, relative to a benchmark (Figure 6). In particular, ALPARC—DEU and—ENG produced a lower PRI than the baseline lexicons and the benchmark for both TP-uniform position-fixed and TP-structured streams. Still, the benchmark outperformed the baseline across TP modes (Table 3). To show that the PRI reduction is consistent across features, we also depict the distributions of PRI for all individual phonological features (Figure S2). For validation, we simulated streams based on ALPARC—DEU lexicons with 2 or 4 syllables per pseudoword and with 3 or 5 pseudowords per lexicon (Figure S3). These results confirm that feature controls consistently reduce PRI across features.

**Figure 6.**
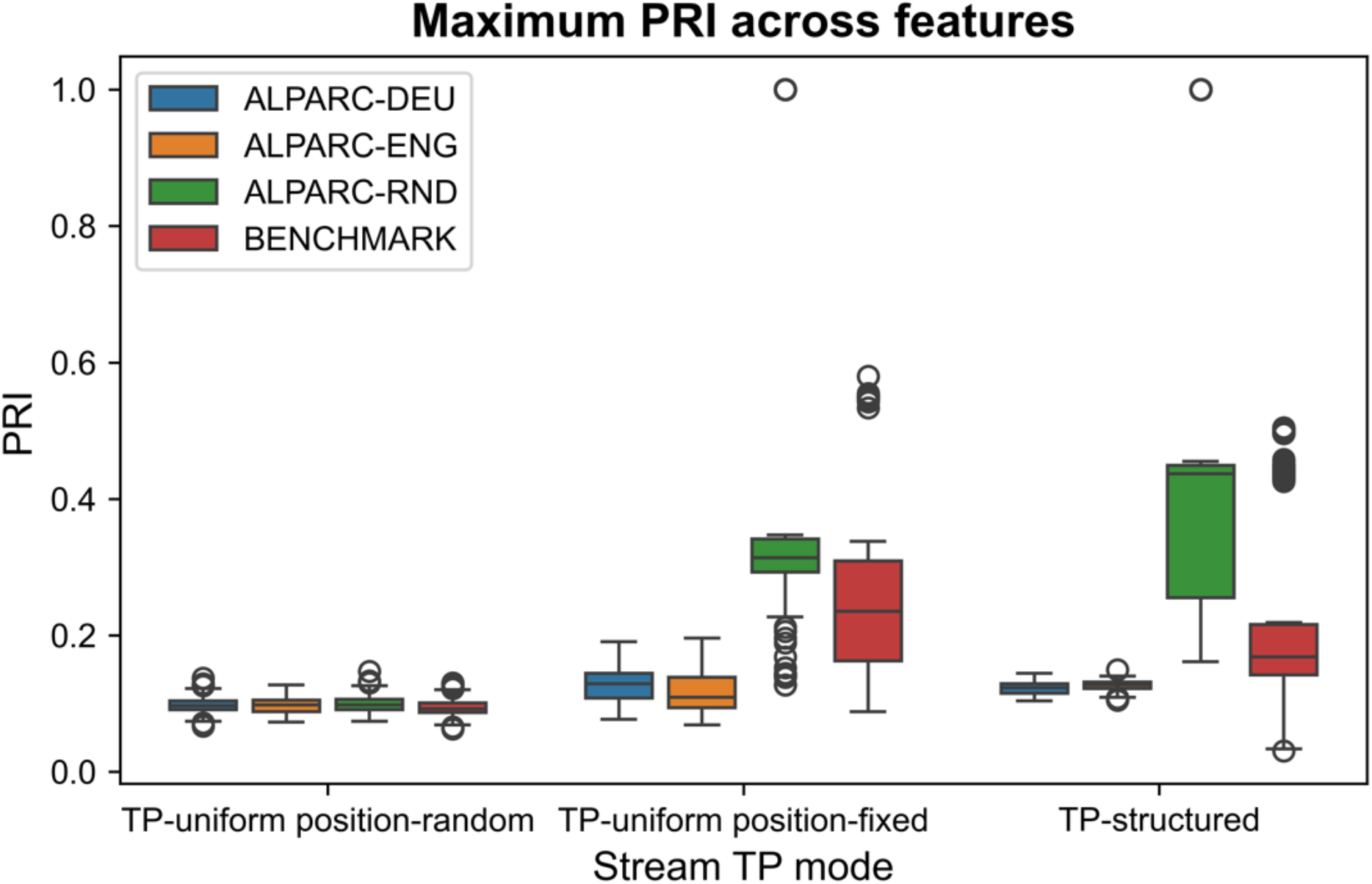
PRI distributions. Boxplots of the distributions of maximum PRI across features for each stream TP mode (n = 10 streams per mode per lexicon) and lexicon type (n = 21 per type). Circles represent outliers.

ALPARC—DEU and —ENG were created as lexicon registers with minimal feature overlap at the rate of pseudowords. This overlap is quantified at the stream level by the PRI. To illustrate that the phonological controls of ALPARC lexicons influence the maximum PRI across features at the stream level, we correlated the feature overlap from 84 lexicons with the maximum PRI. Our results show that controlled lexicons pre-emptively reduce the PRI the stream level, as we found high Pearson’s correlations in TP-uniform position-fixed (*r* = .82) and TP-structured (*r* = .88), between these metrics (Figure S4).

## 5. Discussion

We presented ALPARC as an open-source toolbox to improve stimulus control in linguistic SL studies. We show how ALPARC creates Artificial Languages with Phonological and Acoustic Rhythmicity Controls, which are rendered as TP-uniform streams. ALPARC fulfils three of our aims. First, it crafts artificial lexicons controlling for background statistics based on speech corpora. Second, it generates sequences of tokens with controlled TP distributions in three modalities that contrast empirical surprisal at TP boundaries, factoring out low-level acoustic regularities at the rate of fixed TP patterns, marking pseudowords. Third, it combines these two outputs and computes an index of phonological regularities at the rate of TP boundaries. Our results show that ALPARC improves stimulus controls in artificial languages in terms of lexical, phonological, acoustic, prosodic, and statistical regularities that support SL. Beyond this scope, the three modules of ALPARC may be used independently, and expanded on, to overcome current uncertainties in SL research.

### 5.1. ALPARC—1: The Artificial Language builder

ALPARC—1 provides an interface to moderate the interference of prior linguistic background on the learning of artificial lexicons (Elazar et al., 2022; Siegelman et al., 2018; Varela et al., 2024). To that end, ALPARC— 1 normalizes the frequency distributions of syllables and phoneme co-occurrences in the artificial register, with respect to language-specific corpora statistics. Beyond their relevance for behavioral SL, our linguistic controls might further support neural inference in frequency-tagging studies. While (lexical) frequency and transitional constraints are computationally dissociable (Erickson & Thiessen, 2015), both of these sources of information are tracked by the brain (Tobia et al., 2012). Therefore, both behavioral and neural tracking studies may benefit from language-specific frequency controls oriented to mitigate this confound. Yet, compromises on the extent to which these linguistic controls can be implemented ought to be made on a case-by-case basis to obtain a pseudoword register of adequate size that would allow for further controls (Table S2).

Beyond using corpus data, ALPARC—1 implements phonotactic controls based on OCP constraints. ALPARC—1 considers three phonotactic “rules” as positional repetitions of phonological classes. ALPARC is designed to handle a wide array of features (21) times a very large number of phonemes (5935). From these features, it extracts the manner and place of articulation, avoiding repetitions by lowering their autocorrelation in a sequence at progressively larger lags. OCP-place was previously shown to partially account for neural tracking at the rate of pseudowords (Batterink & Paller, 2017; van der Wulp, 2021). Moreover, phonotactic cues were found to bias speech segmentation behaviorally (Onnis et al., 2005; Onnis & Thiessen, 2013). Building on this evidence, ALPARC offers a systematic way to control the OCP-place, -manner, and -vowel during pseudoword generation, affording phonotactic controls that might be relevant for future SL studies.

ALPARC—1 does not generate lexicon registers made of differently sized units and structures. This design choice was made because an unaware combination of those units is prone to errors that would likely interfere with rhythmicity controls in this implementation. Different phonemic or syllabic structures are not currently supported as they go beyond our scope of producing rhythmic stimuli. Rhythmicity optimization eases further computations in the current execution, in line with the goal of producing rhythmic TP-uniform streams. This way, ALPARC reduces the confounding rhythmicity of phonological features at the rate of TP patterns, making it suitable for study designs that require strict rhythmicity controls (e.g., frequency-tagging). Beyond the current scope, users may also import own data to enter our standard pipeline at any step.

### 5.2. ALPARC—2: The TP chain controller

ALPARC—2 was developed to precisely manipulate TPs in a series of tokens. In doing that, ALPARC replaces a naive randomness approach with a greedy minimum-count rule for increased TP uniformity and precision especially in short sequences. This outcome is most desirable when there are experimental time constraints.

Contrary to classical random-walks, ALPARC—2 sets a uniformity constraint on next-token selection to always explore all possible paths evenly based on a memory matrix. Based on the Fundamental Theorem of Markov Chains, TPs from classical random-walks approach stationarity slowly, as the number of elements in the sequence grows. For instance, naive randomness approaches would pre-shuffle the edge lists randomizing the result. Moreover, a greedy edge selection as a plain Hierholzer sampler satisfies the uniformity condition globally, but may be highly non-uniform early in the stream with a slow convergence to uniform TPs. Naive solutions, such as montecarlo and rejection sampling are suboptimal to the goal of generating short sequences of tokens (< 500 units) with stable TPs. In other words, one is ineffective for short sequences and the other is inefficient for long ones. In fact, the former requires an unknown mixing time to converge to the desired TPs, whereas the latter has an acceptance rate that quickly decays as a function of sequence length. In contrast, ALPARC—2 creates with a stable contrast of surprisal and entropy, after a few samples.

A high TP contrast at pseudoword boundaries is regarded as one key driver of SL. It was proposed that learners detect word boundaries as the locations of high surprisal. This surprisal difference links directly to the computational basis of statistical word learning (Aslin et al., 1998). We defined surprisal as a *stream-level* information metric, in the sense that it is estimated from all transitions in the stream with no knowledge of word structure, and it converges to log_2_(N−1) everywhere. It does not give zero within-word surprisal even in the word-structured condition, because the empirical TP only asymptotically learns that e.g. “ba” is always followed by “go”, after many exposures, never seeing any other continuation. In contrast, a *structure-aware surprisal* is computed under a model that has already inferred word boundaries, whereby TPs are estimated separately within and across words. This gives the theoretically expected values (i.e., 0 within words, and log_2_(W−1) at boundaries). Reporting both information quantities as a function of stream position is informative as the gap between them quantifies how much a listener *gains* by inferring word structure, which is a direct measure of the utility of segmentation and maps onto ideal-observer models of SL.

ALPARC—2 is limited to generating TP-uniform streams of syllables. Therefore, more complex aspects of SL, such as non-adjacent dependency learning, learning of higher hierarchical structures, or entropy-based manipulations, are not within the current scope. However, ALPARC might be considered as a base tool to these ends and its functions might be adapted to specific research purposes in SL. For instance, future work might build on the greedy selection count strategy for sampling from non-uniform distributions, similarly to how we adopted a greedy minimum-count solution to the EC walk. This type of stream generation procedure might produce variable surprisal and entropy at decision chokepoints. While the present implementation was designed to impose TP uniformity through a linearly increasing common cap on the memory transition matrix, ALPARC—2 was designed ad-hoc to have full control on the limits of a memory transition matrix token-by-token. As such, it might serve as a base tool to construct more dynamic TP distributions, by biasing the decision possibilities at the transition points and interfering with the internal constraints of the transition memory matrix between adjacent tokens. Beyond local TPs, filler syllables afford studying non-adjacent dependency learning.

ALPARC—2 improves construct validity at the stimulus level to strengthen neural inference. Yet, it is important to note that neural peaks at the pseudoword rate do not uniquely index TP computation. In fact, such peaks may also reflect other segmentation mechanisms, such as repetition-based learning, positional regularity learning, or surprisal dynamics based on learned multi-syllabic groupings. While ALPARC controls surprisal dynamics of short-term local TPs, the neural peaks at the pseudoword rate may also result from other long-term memory-based grouping mechanisms (Jessop et al., 2025).

ALPARC—2 includes audio editing tools to obtain frequency-tagged streams of syllables. Individually synthesized (or naturally recorded) syllables can be concatenated according to the sequences of ALPARC lexicon and streams. In the current implementation, our audio editing scripts allow for equalization of syllable duration and, amplitude, and manipulations of pitch tiers to imitate periodic intonational prosodic boundaries (see also Lamekina & Meyer, 2023 for details). While this feature is optimal for frequency-tagging factorial designs to disentangle statistical and prosodic influences on linguistic SL in rhythmic, these stimuli have a low acoustic realism, which might be raised by natural voice recordings (Alexandrou et al., 2020).

To gain acoustic realism, users may adopt their own audio synthesis or recording strategies. While all of ALPARC features can in principle generalize to human recorded speech, ALPARC disregards coarticulation during lexicon generation to facilitate downstream rhythmicity controls and to avoid the manual cutting of the audio files at coarticulated syllabic boundaries. With the same limitation, ALPARC affords changing the pitch of a (synthesized) audio to a natural strong-weak-weak contour or a synthesis of hundreds of German prosodic contours (see Lamekina & Meyer, 2023 for details). Beyond audio editing for frequency-tagging, ALPARC handles sequences of syllables that do not necessarily need to be presented periodically. For instance, a parsed phonemic transcript is compatible with other ALPARC functionalities.

Beyond providing additional support tools for audio editing, it is important to note that ALPARC—2 generates uniform TP chains as sequences of symbolic digits. As such, this module is not linguistic per se and offers low-level positional controls that guarantee stable surprisal values in any sort of statistical stream. Thus, the strategy that ALPARC—2 uses for making TP chains may be useful for SL research across domains.

ALPARC—2 supports the use of a TP-uniform baseline, matching TP-structured positional constraints. ALPARC applies an artificial language to a TP-uniform stream, thereby mapping each token to a pseudoword (in TP-structured streams) or to a syllable (in TP-uniform streams), while expanding TP-structured streams of pseudowords into connected streams of syllables. In the linguistic domain, syllable positional constraints have been shown to induce systematic acoustic differences that are reflected in the amplitude modulation spectra (Pinto et al., 2022). ALPARC streams are in line with this finding, showing that spectral profiles of the TP-uniform position-random baseline does not reflect the same peak at the repetition rate that are present in TP-structured streams, whereas a TP-uniform position-fixed does equalize acoustic regularities (Pinto et al., 2022).

### 5.3. ALPARC—3: The rhythmicity checker

ALPARC—3 diagnoses artificial language stimuli to visualize their PRI distributions. Our results show that ALPARC lexicons reduce the maximum PRI relative to random lexicons of pseudowords that disregard feature controls, but also to benchmarking lexicons used in prior studies on linguistic SL. While the latter certainly attained a better phonological rhythmicity control than our baseline, which must be due to the expert conduct of the prior studies, ALPARC outperformed this benchmark. Lower PRIs denote a lower impact that the periodicity of phonological features may impose on the syllable streams. This confound is further eased if corresponding TP-structured and TP-uniform streams have the same PRI. Lexicons and streams generated with ALPARC may thus allow for assessing SL of rhythmic TP patterns matching out phonological regularities in TP-structured and position-controlled baseline streams. Future work might expand on this module by mapping the exact stream location of an impulse along the vector, rather than returning only an average of impulses across all stream locations. This output might be more informative to study dynamic phonological patterns.

The PRI might be useful in future SL studies to assess neural tracking of phonological features. For instance, minimal edits to the function that generates the artificial lexicons could lead to the creation of speech streams—that maximize the PRI of one, or more, features, in order to study the neural tracking of specific phonological features systematically. In this broader scope, ALPARC might support the study of phonemic mapping through pattern repetitions, given the arbitrary size and shape of the binary matrix and kernels.

ALPARC—3 can be useful for meta-analyses purposes. The PRI is a metric that diagnoses instances of phonological regularities at the TP rate in a syllable stream. ALPARC considers syllables as basic acoustic, phonological, and statistical units. Lexicons with different numbers of syllables per word are not supported by ALPARC—1 during generation. Yet, ALPARC—3 can diagnose these lexicons, as it returns a PRI from such streams after mapping the locations of any desired kernel (even non-uniformly periodic, e.g., [1 1 0 1 0 1]) in feature space. The convolution of the TP kernels spans the whole matrix made of syllables unfolding at a fixed rate (columns) and the corresponding feature vectors of the phonemes of the syllables (rows). The patterning of the TP kernels and the feature matrix can be set by the user to depends on particular syllable’s phonemes (e.g., our default settings consider different features for consonants [C] and vowels [V]). In our pipeline, we limit the syllable structure to CV syllables to yield a size-compatible matrix of phonological features across syllables. Still, if CVC-CV words (even with coarticulation) are to be used, and their exact IPA transcription is found in the list, it should be possible to build a custom phonological vector of the syllable by stacking the phonological features that are considered as relevant for each phoneme and padding the shortest stack with zeros (e.g., for syllables that do not present that features). This nominal consistency will result in interpretable PRIs in CVC-CV, but not in a CVC-VCV word, as in our incorporation of the benchmark lexicons.

### 5.4. Conclusion

We presented ALPARC to generate and diagnose artificial languages with ad-hoc rhythmicity controls. We summarize three main functionalities. First, ALPARC crafts highly controlled lexicon registers through filtering linguistic corpora statistics. Second, ALPARC crafts uniform TP sequences in three critical contrasts, with one baseline that equalizes low-level (acoustic) confounds and another that maximizes the information difference at the salient boundaries. These modules combine to make streams that contrast TPs at syllable or pseudowords boundaries. Third, ALPARC computes a PRI on the phonological feature matrix that reflects the stream of syllables, quantifying the density of (rhythmic) binary kernels. Our results show that ALPARC succeeds in the intended purpose of generating frequency-tagged audios that contrast prosodic and statistical regularities while factoring out some of the most noticeable linguistic, acoustic, and phonological confounds. Our results show that not controlling for these factors might result in differences at baseline. To date, ALPARC produces stimuli for this purpose in less than a minute. While ALPARC is currently optimized for frequency-tagging designs, we specify how its three modules may not be fully limited to this specific paradigm and might independently serve as base tools to meet specific research purposes in SL research across domains.

## Supporting information

alparc_supplement

## Acknowledgments

The study was supported by the Max Planck Society through the award of the Independent Max Planck Research Group Language Cycles to LM. NM was supported by BMBF (Federal Ministry of Education and Research) through ACONITE (01IS22065), and the Center for Scalable Data Analytics and Artificial Intelligence (ScaDS.AI.) Leipzig, and the Max Planck IMPRS CoNI Doctoral Program. The authors would like to thank Dimitra Kiakou and Shashank Shetty Kalavara for early insights into the implementation of TP controls in the early stages of development, Yulia Lamekina for insights on prosody manipulations, and Jule Nabrotzky for comments on the manuscript. We would also like to thank the Bogaerts Lab for early insightful comments into potential future developments. LT and LM conceptualized the research. LT, NM, and LM conducted the formal analyses. LT and NM developed the software. LT and NM created the visualizations. LM supervised the research. LM acquired funding. LT wrote the manuscript. LT, NM, and LM edited the manuscript. The authors declare no competing interests.

