## Supplementary material for "ALPARC: Artificial Languages with Phonological and Acoustic Rhythmicity Controls": alparc_supplement

### Supplementary Information for ALPARC: Artificial Languages with Phonological and Acoustic Rhythmicity Controls

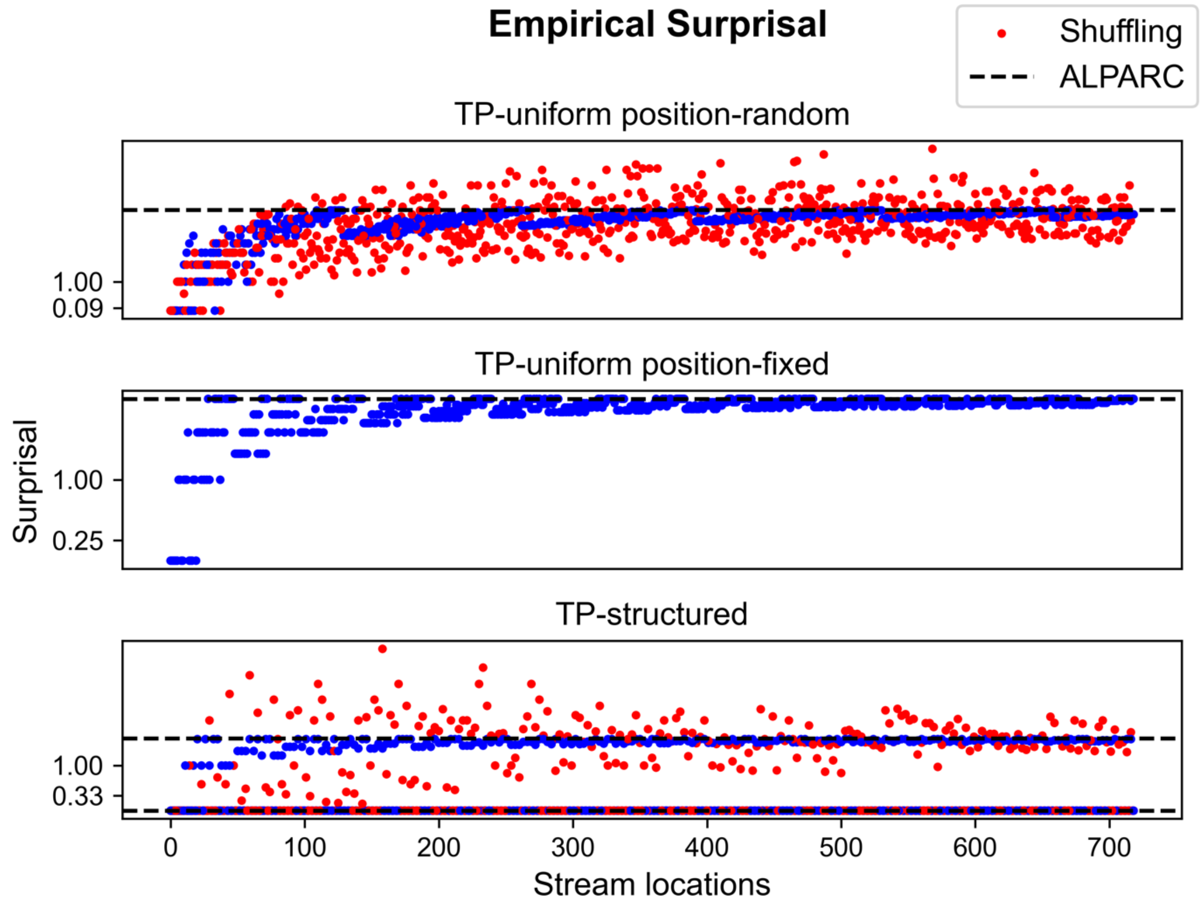


**Figure S1. Empirical surprisal.** Surprisal (bits) is plotted as a function of stream location contrasting a shuffling method (red dots) and ALPARC streams (blue dots). The horizontal black line indicates the theoretical surprisal at TP boundary.

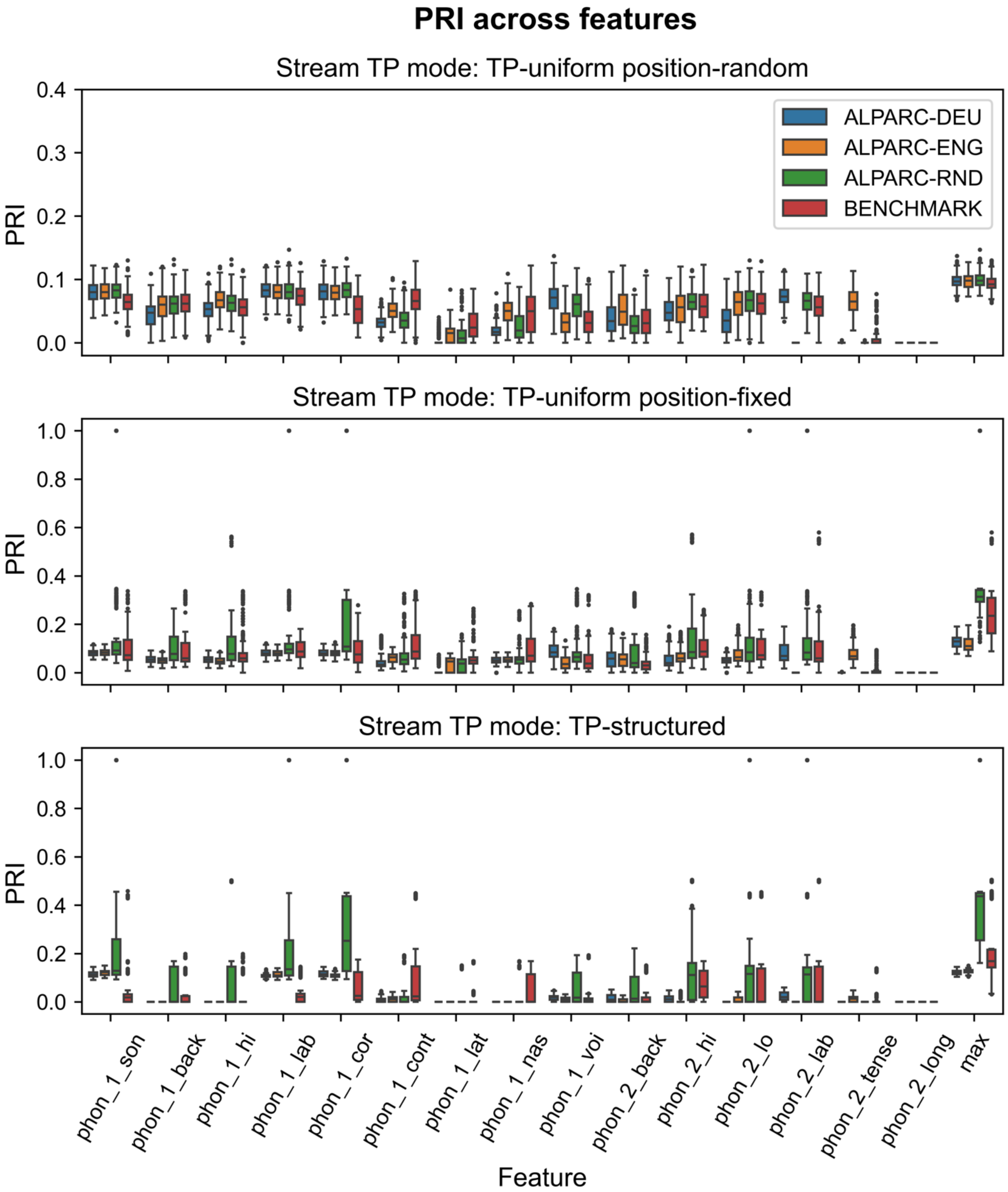


**Figure S2. Distributions of PRI for each feature.** The PRI is plotted as a function of individual phonological features (abscissa), lexicon type (colored boxes) and stream TP modes (separate rows). Circles represent outliers. Consonant- and vowel-relevant features are referred to as "phon_1" and "phon_2", respectively. Other abbreviations as in the main text.

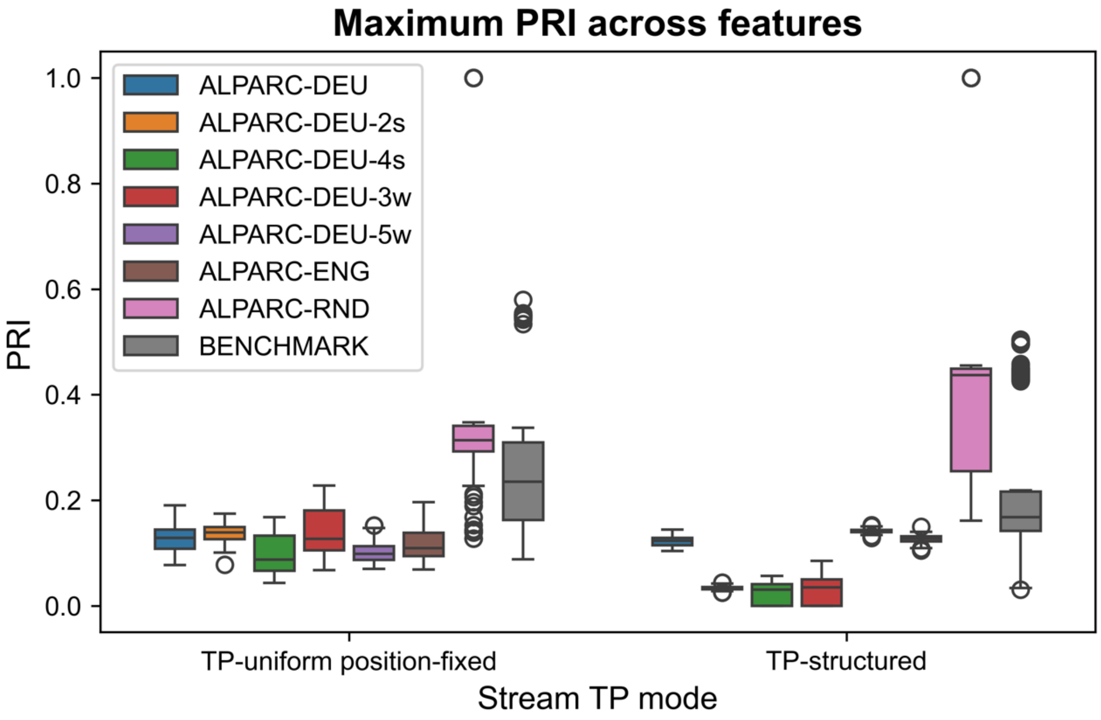


**Figure S3. PRI distributions for lexicons with different number of syllables and pseudowords.** Boxplots of the distribution of maximum PRI across features as a function of stream TP mode (abscissae) and lexicon type (from left to right, color-coded in the legend). The number of syllables per pseudoword and the number of pseudowords per lexicon are abbreviated by the number preceding the letter "s" and "w", respectively, only if these values differ from the default (s = 3; w = 4). Abbreviations for lexicon type as in Figure 4. Circles represent outliers.

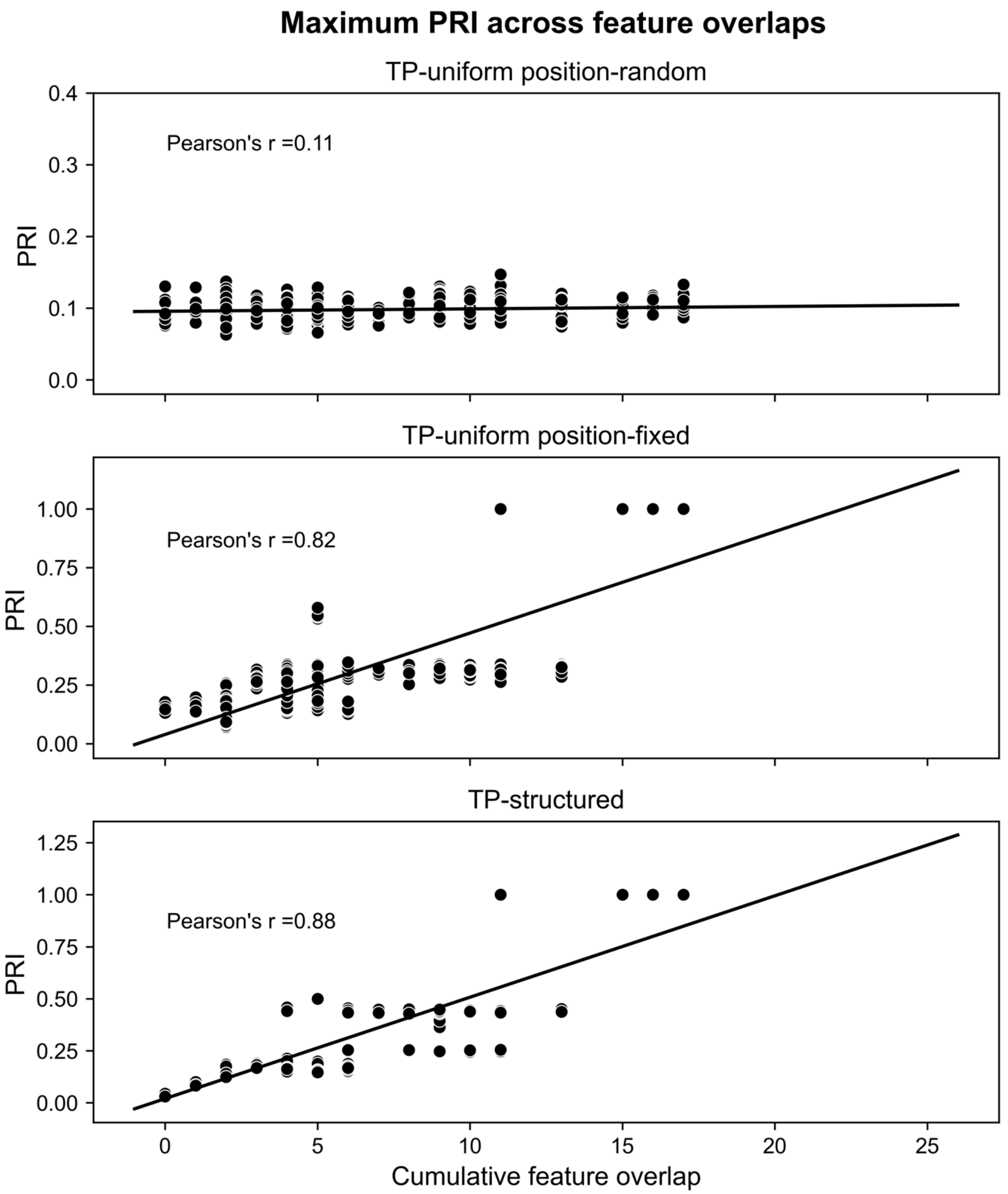


**Figure S4. Feature overlap correlates with phonological rhythmicity index (PRI).** The maximum PRI is plotted as a function of cumulative feature overlap for 84 artificial lexicons. Every dot represents the average maximum PRI across 10 streams per TP mode for each lexicon. The interpolating line represents the linear fit to the data. Pearson's r for each stream TP mode (separate rows) indicate the correlation coefficients between the cumulative feature overlap computed at the lexicon level, and the maximum PRI from those lexicons averaged across 10 streams.

**Table S1. Benchmark Lexicons.**

| Lexicon | Word 1 | Word 2 | Word 3 | Word 4 | Reference |
|---|---|---|---|---|---|
| 1 | pi.ɾu.ta | ba.go.li | to.ku.da | gu.ki.bo | (Cunillera et al., 2009) |
| 2 | pa.be.la | di.ne.ka | lu.fa.ri | xi.so.du | (Cunillera et al., 2009) |
| 3 | ma.xu.pe | xe.ro.ga | de.mu.si | fo.le.ti | (Cunillera et al., 2009) |
| 4 | pu.ke.mi | ra.fi.nu | bi.na.po | me.do.xi | (Cunillera et al., 2009) |
| 5 | no.ni.xe | bu.lo.te | re.mo.fu | ko.tu.sa | (Cunillera et al., 2009) |
| 6 | mi.lo.de | da.le.bu | no.ru.pa | ka.te.xi | (Cunillera et al., 2006) |
| 7 | ne.do.li | ri.fo.nu | ba.to.gu | ki.ra.pu | (Cunillera et al., 2006) |
| 8 | go.na.be | mu.di.la | ro.ni.xe | pi.ku.sa | (Cunillera et al., 2006) |
| 9 | fu.bi.re | xe.tu.si | ta.fi.ko | ke.ma.po | (Cunillera et al., 2006) |
| 10 | ti.fa.xu | so.du.xi | me.lu.bo | ga.ni.pe | (Cunillera et al., 2006) |
| 11 | mi.po.la | za.bɛ.tu | ʁo.ki.sɛ | nu.ga.di | (Kiai & Melloni, 2021) |
| 12 | dɛ.mʊ.ri | sɛ.ni.gɛ | ræ.ku.səʊ | pi.lɛ.ru | (Assaneo et al., 2019) |
| 13 | ki.fəʊ.bu | lu.fɑ.gi | pæ.beɪ.lɑ | tɑ.gəʊ.fʊ | (Assaneo et al., 2019) |
| 14 | bi.du.pɛ | məʊ.bɑ.li | rɛ.gæ.tʊ | sæ.tɛ.kəʊ | (Assaneo et al., 2019) |
| 15 | bəʊ.dɑ.mɛ | fi.nəʊ.pɑ | gʊ.rɑ.təʊ | ləʊ.kæ.neɪ | (Assaneo et al., 2019) |
| 16 | fɛ.si.nɑ | kɛ.su.dəʊ | mæ.pʊ.di | ti.mi.nu | (Assaneo et al., 2019) |
| 17 | tu.pi.ɹoʊ | goʊ.la.bu | pa.doʊ.ti | bi.da.ku | (Batterink & Paller, 2017) |
| 18 | meɪ.lu.gi | ɹa.fi.nu | pu.keɪ.mi | toʊ.na.poʊ | (Batterink & Paller, 2017) |

| Lexicon | Word 1 | Word 2 | Word 3 | Word 4 | Reference |
|---|---|---|---|---|---|
| 19 | goʊ.la.tu | da.ɹoʊ.pi | ti.bu.doʊ | pa.bi.ku | (McNealy et al., 2006) |
| 20 | poʊ.fi.mu | noʊ.vu.ka | vi.koʊ.ga | ba.fu.gi | (McNealy et al., 2006) |
| 21 | ma.nu.toʊ | ni.moʊ.lu | voʊ.ɹi.fa | li.du.ɹa | (McNealy et al., 2006) |

**Table S2. Working example.**

| Lexicon | Language | Phonemes | Syllable fitler α | Bigram (α = none) | Syllables | Words |
|---|---|---|---|---|---|---|
| ALPARC-DEU | deu | cV | .05 | True | 3 | 4 |
| ALPARC-ENG | eng | cv | .05 | True | 3 | 4 |
| ALPARC-RND | deu | cV | .05 | False | 3 | 4 |
| ALPARC-DEU-3w | deu | cV | .05 | True | 3 | 3 |
| ALPARC-DEU-5w | deu | cV | .05 | True | 3 | 5 |
| ALPARC-DEU-2s | deu | cV | .05 | True | 2 | 4 |
| ALPARC-DEU-4s | deu | cV | .05 | True | 4 | 4 |
